# Genome variation and population structure among 1,142 mosquitoes of the African malaria vector species *Anopheles gambiae* and *Anopheles coluzzii*

**DOI:** 10.1101/864314

**Authors:** The Anopheles gambiae 1000 Genomes Consortium, Chris S Clarkson, Alistair Miles, Nicholas J Harding, Eric R Lucas, C J Battey, Jorge Edouardo Amaya-Romero, Jorge Cano, Abdoulaye Diabate, Edi Constant, Davis C Nwakanma, Musa Jawara, John Essandoh, Joao Dinis, Gilbert Le Goff, Vincent Robert, Arlete D Troco, Carlo Costantini, Kyanne R Rohatgi, Nohal Elissa, Boubacar Coulibaly, Janet Midega, Charles Mbogo, Henry D Mawejje, Jim Stalker, Kirk A Rockett, Eleanor Drury, Daniel Mead, Anna E Jeffreys, Christina Hubbart, Kate Rowlands, Alison T Isaacs, Dushyanth Jyothi, Cinzia Malangone, Maryam Kamali, Christa Henrichs, Victoria Simpson, Diego Ayala, Nora J Besansky, Austin Burt, Beniamino Caputo, Alessandra della Torre, Michael Fontaine, H. Charles J Godfray, Matthew W Hahn, Andrew D Kern, Mara K N Lawniczak, Samantha O’Loughlin, Joao Pinto, Michelle M Riehle, Igor Sharakhov, Daniel R Schrider, Kenneth D Vernick, Bradley J White, Martin J Donnelly, Dominic P Kwiatkowski

## Abstract

Mosquito control remains a central pillar of efforts to reduce malaria burden in sub-Saharan Africa. However, insecticide resistance is entrenched in malaria vector populations, and countries with high malaria burden face a daunting challenge to sustain malaria control with a limited set of surveillance and intervention tools. Here we report on the second phase of a project to build an open resource of high quality data on genome variation among natural populations of the major African malaria vector species *Anopheles gambiae* and *Anopheles coluzzii*. We analysed whole genomes of 1,142 individual mosquitoes sampled from the wild in 13 African countries, and a further 234 individuals comprising parents and progeny of 11 lab crosses. The data resource includes high confidence single nucleotide polymorphism (SNP) calls at 57 million variable sites, genome-wide copy number variation (CNV) calls, and haplotypes phased at biallelic SNPs. We used these data to analyse genetic population structure, and characterise genetic diversity within and between populations. We also illustrate the utility of these data by investigating species differences in isolation by distance, genetic variation within proposed gene drive target sequences, and patterns of resistance to pyrethroid insecticides. This data resource provides a foundation for developing new operational systems for molecular surveillance, and for accelerating research and development of new vector control tools.

## Introduction

The 10 countries with the highest malaria burden in Africa account for 65% of all malaria cases globally, and attempts to reduce that burden further are stalling in the face of significant challenges [1]. Not least among these, resistance to pyrethroid insecticides is widespread throughout African malaria mosquito populations, potentially compromising the efficacy of mosquito control interventions which remain a core tenet of global malaria strategy [2, 3]. There is a broad consensus that further progress cannot be made if interventions are applied blindly, but must instead be guided by data from epidemiological and entomological surveillance [4]. Genome sequencing technologies are considered to be a key component of future malaria surveillance systems, providing insights into evolutionary and demographic events in mosquito and parasite populations that are otherwise difficult to obtain [5]. Genomic surveillance systems will not work in isolation, but will depend on high quality open genomic data resources, including baseline data on genome variation from multiple mosquito species and geographical locations, against which comparisons can be made and inferences regarding new events can be drawn.

Better surveillance can increase the impact and longevity of available mosquito control tools, but sustaining malaria control will also require the development and deployment of new tools [4]. This includes repurposing existing insecticides not previously used in public health [6, 7], developing entirely new insecticide classes, and developing tools that don’t rely on insecticides, such as genetic modification of mosquito populations [8]. Research and development of new mosquito control tools has been greatly facilitated by the availability of open genomic data resources, including high quality genome assemblies [9, 10], annotations [11], and more recently by high quality resources on genetic variation among natural mosquito populations [12]. Further expansion of these open data resources to incorporate unsampled mosquito populations and new types of genetic variation can provide new insights into a range of biological and ecological processes, and help to accelerate scientific discovery from basic biology through to operational research.

The *Anopheles gambiae* 1000 Genomes (Ag1000G) project^1^ was established in 2013 to build a large scale open data resource on natural genetic variation in malaria mosquito populations. The Ag1000G project forms part of the Malaria Genomic Epidemiology Network^2^ (MalariaGEN), a data-sharing community of researchers investigating how genetic variation in humans, mosquitoes and malaria parasites can inform the biology, epidemiology and control of malaria. The first phase of the Ag1000G project released data from whole genome Illumina deep sequencing of the major Afrotropical malaria vector species *Anopheles gambiae* and *Anopheles coluzzii* [12], two closely related siblings within the *Anopheles gambiae species complex* [13]. Mosquitoes were sampled in 8 African countries from a broad geographical range, spanning Guinea-Bissau in West Africa to Kenya in East Africa. Genetic diversity was found to be high in most populations, but there were marked patterns of population structure, and clear differences between populations in the magnitude and architecture of genetic diversity, indicating complex and varied demographic histories. However, both of these species have a large geographical range [14], and many countries and ecological settings are not represented in the Ag1000G phase 1 resource. Also, only SNPs were studied in Ag1000G phase 1, but other types of genetic variation are known to be important. In particular, copy number variation has long been suspected to play a key role in insecticide resistance [15, 16, 17], but no previous attempts to call genome-wide CNVs have been made in these species.

This paper describes the data resource produced by the second phase of the Ag1000G project. Within this phase, sampling and sequencing was expanded to include additional wild-caught mosquitoes collected from five countries not represented in phase 1. This includes three new locations with *An. coluzzii*, providing greater power for genetic comparisons with *An. gambiae*, and two island populations, providing a useful reference point to compare against mainland populations. Seven new lab crosses are also included, providing a substantial resource for studying genome variation and recombination within known pedigrees. In this phase we studied both SNPs and CNVs, and rebuilt a haplotype reference panel using all wild-caught specimens. Here we describe the data resource, and use it to re-evaluate major population divisions and characterise genetic diversity. We also illustrate the broad utility of the data by comparing geographical population structure between the two mosquito species to investigate evidence for differences in dispersal behaviour; analyse genetic diversity within a gene in the sex-determination pathway currently targeted for gene drive development; and provide some preliminary insights into the prevalence of different molecular mechanisms of pyrethroid resistance.

## Results

### Population sampling and sequencing

We performed whole genome sequencing of 377 individual wild-caught mosquitoes, including individuals collected from 3 countries (The Gambia, Côte d’Ivoire, Ghana) and two oceanic islands (Bioko, Mayotte) not represented in the previous project phase. We also sequenced 152 individuals comprising parents and progeny from seven lab crosses, where parents were drawn from the Ghana, Kisumu, Pimperena, Mali and Akron colonies. We then combined these data with the sequencing data previously generated during phase 1 of the project, to create a total resource of data from 1,142 wild-caught mosquitoes (1,058 female, 84 male) from 13 countries (Figure 1; Table S1) and 234 mosquitoes from 11 lab crosses (Table S2). As in the previous project phase, all mosquitoes were sequenced individually on Illumina technology using 100 bp paired-end reads to a target depth of 30X, and all 1,142 mosquitoes in the final resource had a mean depth above 14X.

**Figure 1.**
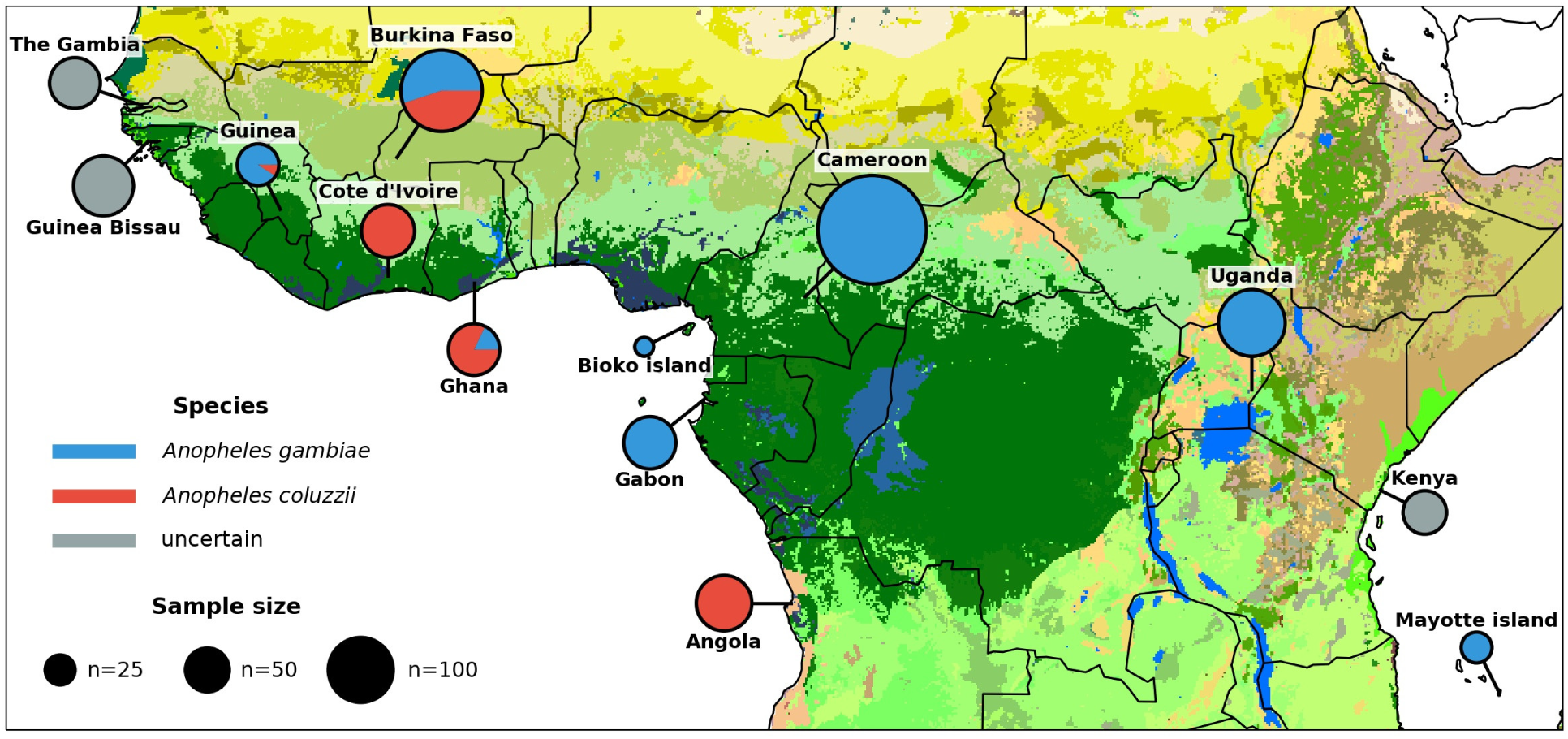
Ag1000G phase 2 sampling locations. Colour of circle denotes species and area represents sample size. Species assignment is labelled as uncertain for samples from Guinea-Bissau, The Gambia and Kenya, because all individuals from those locations carry a mixture of *An. gambiae* and *An. coluzzii* ancestry informative markers, see main text and Figure S1 for details. Map colours represent ecosystem classes, dark green designates forest ecosystems; see Figure 9 in [18] for a compete colour legend.

### Genome variation

Sequence reads from all individuals were aligned to the AgamP3 reference genome [9, 10] and SNPs were discovered using methods described previously [12]. In total, we discovered 57,837,885 SNPs passing all variant quality filters, 11% of which were newly discovered in this project phase. Of these high quality SNPs, 24% were found to be multiallelic (three or more alleles). We also analysed genome accessibility to identify all genomic positions where read alignments were of sufficient quality and consistency to support accurate discovery and genotyping of nucleotide variation. Similar to the previous project phase, we found that 61% (140 Mbp) of genome positions were accessible, including 91% (18 Mbp) of the exome and 58% (121 Mbp) of non-coding positions. Overall we discovered an average of one variant allele every 1.9 bases of the accessible genome. We then used high quality biallelic SNPs to construct a new haplotype reference panel including all 1,142 wild-caught individuals, via a combination of read-backed phasing and statistical phasing as described previously [12].

In this project phase we also performed a genome-wide CNV analysis, described in detail elsewhere [19]. In brief, for each individual mosquito, we called CNVs by fitting a hidden Markov model to windowed data on depth of sequence read coverage, then compared calls between individuals to identify shared CNVs. The CNV callset comprises 31,335 distinct CNVs, of which 7,086 were found in more than one individual, and 1,557 were present at at least 5% frequency in one or more populations. CNVs spanned more than 68 Mbp in total and overlapped 7,190 genes. CNVs were significantly enriched in gene families associated with metabolic resistance to insecticides, with three loci in particular (two clusters of cytochrome P450 genes *Cyp6p/aa*, *Cyp9k1* and a cluster of glutathione S-transferase genes *Gste*) having a large number of distinct CNV alleles, multiple alleles at high population frequency, and evidence that CNVs are under positive selection [19]. CNVs at these loci are thus likely to be playing an important role in adaptation to mosquito control interventions.

### Species assignment

The conventional and most widely used molecular assays for differentiating *An. gambiae* from *An. coluzzii* are based on fixed differences in the centromeric region of the X chromosome [20, 21]. In the first phase of the Ag1000G project, we compared the results of these assays with genotypes at 506 ancestry-informative SNPs distributed across all chromosome arms, and found that in some cases the conventional assays were not concordant with species ancestry at other genome locations. In particular, all individuals from two sampling locations (Kenya, Guinea-Bissau) carried a mixture of *An. gambiae* and *An. coluzzii* alleles, creating uncertainty regarding the appropriate species assignment [12]. Applying the same analysis to the new samples in Ag1000G phase 2, we found that mosquitoes from The Gambia also carried a mixture of alleles from both species, in similar proportions to mosquitoes from Guinea-Bissau (Figure S1). In all other locations, alleles at ancestry-informative SNPs were concordant with conventional diagnostics [20, 21], except on chromosome arm 2L where there has been a known introgression event carrying an insecticide resistance allele from *An. gambiae* into *An. coluzzii* [22, 23, 24, 25]. We observed this introgression in *An. coluzzii* from both Burkina Faso and Angola in the phase 1 cohort, and it was also present among *An. coluzzii* from Côte d’Ivoire, Ghana and Guinea in the phase 2 cohort.

### Population structure

We investigated genetic population structure within the cohort of wild-caught mosquitoes by performing dimensionality reduction analyses on the genome variation data, including UMAP [26] and PCA [27] of biallelic SNPs from euchromatic regions of Chromosome 3 (Figure 2; Figure S2), and PCA of CNVs from the whole genome (Figure S3). To complement these analyses, we fitted models of population structure and admixture [28] to the SNP data (Figure S4). We also used SNPs to compute two measures of genetic differentiation, average *F_ST_* and rates of rare variant sharing, between pairs of populations defined by country of origin and species (Figure 3). From these analyses, three major groupings of individuals from multiple countries were evident: *An. coluzzii* from West Africa (Burkina Faso, Ghana, Côte d’Ivoire, Guinea); *An. gambiae* from West, Central and near-East Africa (Burkina Faso, Ghana, Guinea, Cameroon, Uganda); individuals with uncertain species status from far-West Africa (Guinea-Bissau, The Gambia). Within each of these groupings, samples clustered closely in all PCA and UMAP components and in admixture models for up to *K* = 5 ancestral populations, and differentiation between countries was weak, consistent with relatively unrestricted gene flow between countries. Each of the remaining PCA clusters comprised samples from a single country and species (Angola *An. coluzzii*; Gabon *An. gambiae*, Mayotte *An. gambiae*; Bioko *An. gambiae*; individuals with uncertain species status from Kenya), and in general each of these populations was more strongly differentiated from all other populations, consistent with a role for geographical factors limiting gene flow. The admixture analyses for Mayotte and Kenya modelled individuals from both populations as a mixture of multiple ancestral populations. This could represent some true admixture in these populations’ histories, but could also be an artefact due to strong genetic drift [29], and requires further investigation. A comparison of the two *An. gambiae* island populations is interesting because Mayotte was highly differentiated from all other populations, but individuals from Bioko were more closely related to other West African *An. gambiae*, suggesting that Bioko may not be isolated from continental populations despite a physical separation of more than 30 km.

**Figure 2.**
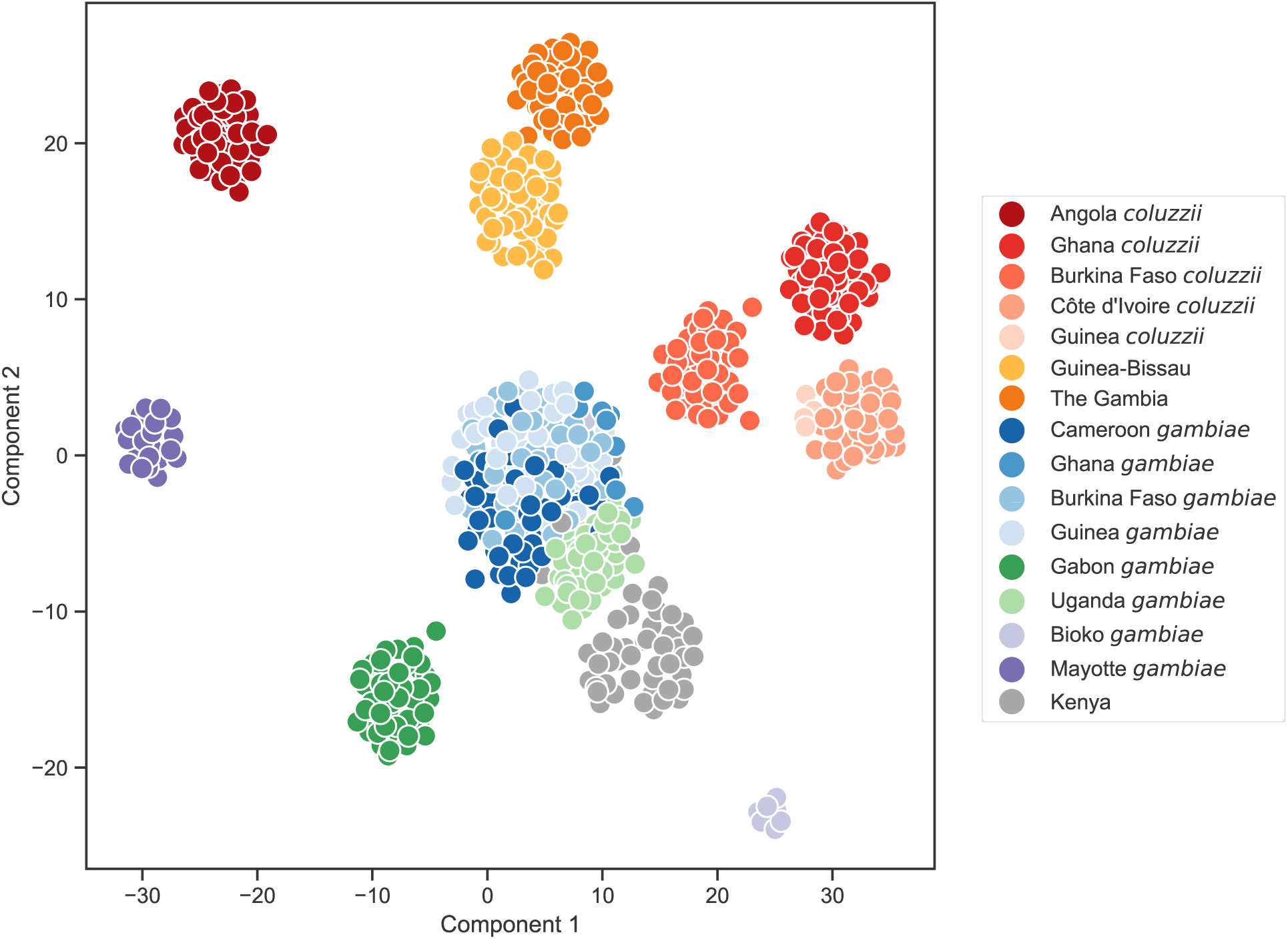
Population structure analysis of the wild-caught mosquitoes using UMAP [26]. Geno-type data at biallelic SNPs from euchromatic regions of Chromosome 3 were projected onto two components. Each marker represents an individual mosquito. Mosquitoes from each country and species were randomly downsampled to at most 50 individuals.

**Figure 3.**
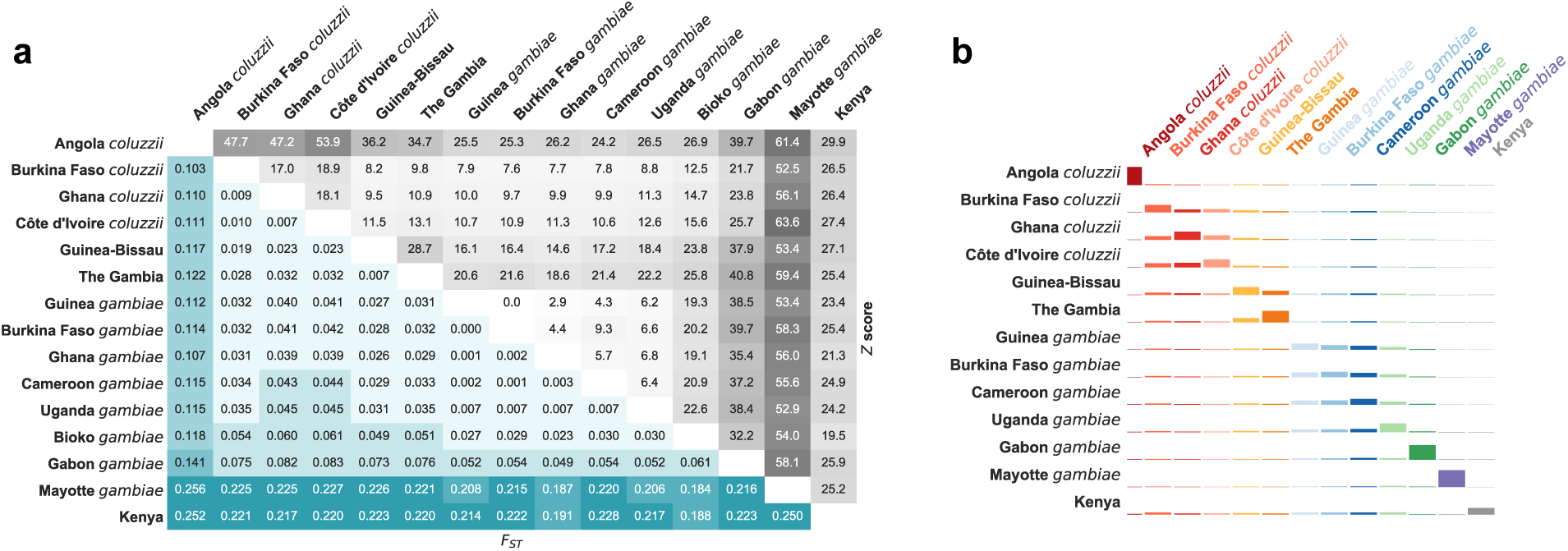
Genetic differentiation between populations, computed using using biallelic SNPs from euchromatic regions of Chromosome 3. **(a)** Average allele frequency differentiation (*F_ST_*) between pairs of populations. The bottom left triangle shows average *F_ST_* values between each population pair. The top right triangle shows the Z score for each *F_ST_* value estimated via a block-jackknife procedure. **(b)** Allele sharing in doubleton (*f*_2_) variants. For each population, we identified the set of doubletons with at least one allele originating from an individual in that population. We then computed the fraction of those doubletons shared with each other population and the fraction shared only within itself. The height of the coloured bars represent the probability of sharing a doubleton allele between or within populations. Heights are normalized row-wise for each population so that the sum of coloured bars in each row equals 1.

The new locations sampled in this project phase allow more comparisons to be made between *An. gambiae* and *An. coluzzii*, and there are many open questions regarding their behaviour, ecology and evolutionary history. For example, it would be valuable to know whether there are any differences in long-range dispersal behaviour between the two species [30] as have been suggested by recent studies in Sahelian regions [31, 32]. Providing a comprehensive answer to this question is beyond the scope of this study, but we performed a preliminary analysis by estimating Wright’s neighbourhood size for each species [33]. This statistic is an approximation for the effective number of potential mates for an individual, and can be viewed as a measurement of how genetic differentiation between populations correlates with the geographical distance between them (isolation by distance). We used Rousset’s method for estimating neighbourhood size based on a regression of normalised *F_ST_* against the logarithm of geographical distance [34]. To avoid any confounding effect of major ecological discontinuities, we used only populations from West Africa and Central Africa north of the equatorial rainforest. We found that average neighbourhood sizes are significantly lower in *An. coluzzii* than in *An. gambiae* (Wilcoxon, *W* = 1320, *P <* 2.2*e −* 16) (Figure 4), indicating stronger isolation by distance among *An. coluzzii* populations and suggesting a lower rate and/or range of dispersal. However, we do not have representation of both species at all sampling locations, and so further sampling will be needed to confirm this result.

**Figure 4.**
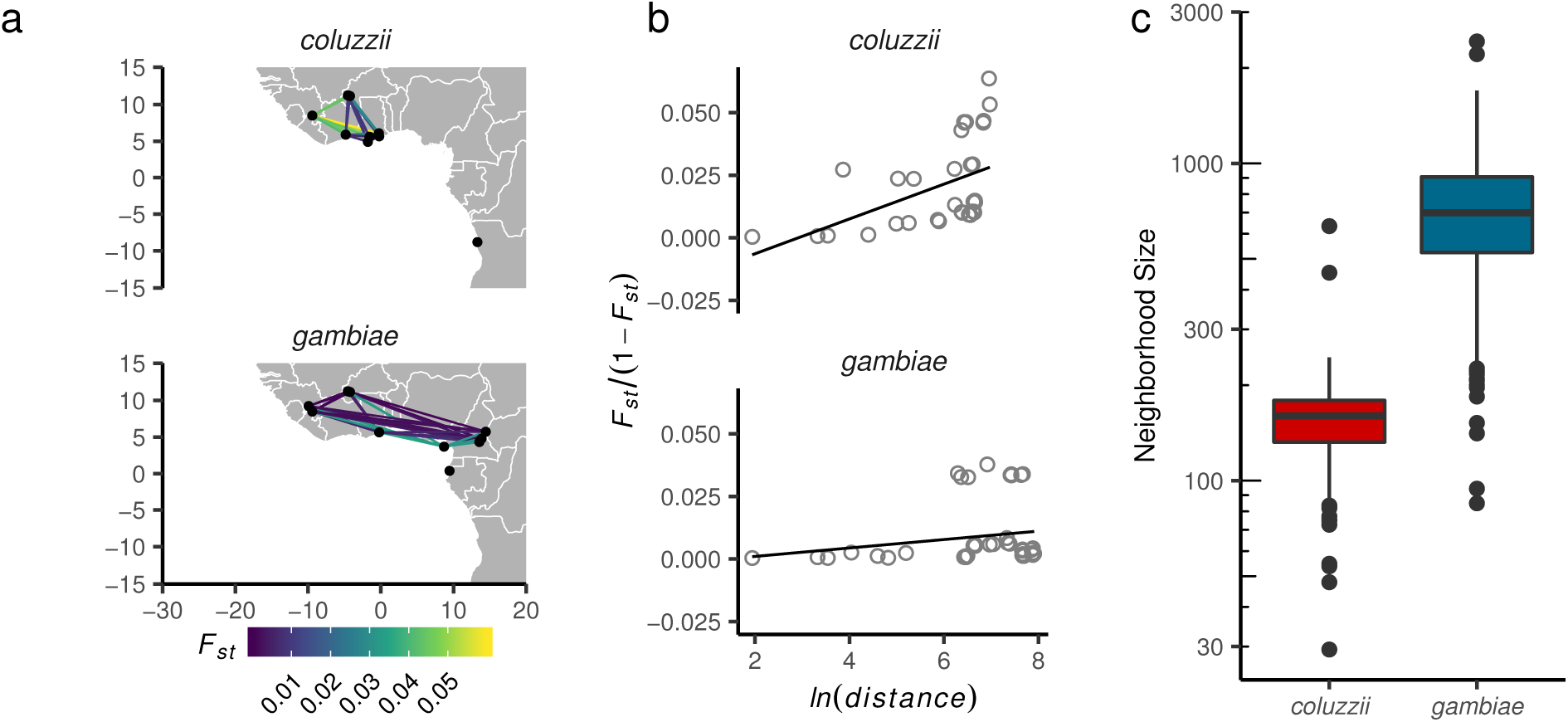
Comparison of isolation by distance between *An. coluzzii* and *An. gambiae* populations from locations in West and Central Africa north of the equatorial rainforest. **(a)** Study region and pairwise *F_ST_*. **(b)** Regressions of average genome-wide *F_ST_* against geographic distance, following Rousset [34]. Neighbourhood size is estimated as the inverse slope of the regression line. **(c)** Difference in neighbourhood size estimates by species. Box plots show medians and 95% confidence intervals of the distribution of estimates calculated in 200 kbp windows across the euchromatic regions of Chromosome 3.

### Genetic diversity

The populations represented in the Ag1000G phase 2 cohort can serve as a reference point for comparisons with populations sampled by other studies at other times and locations. To facilitate population comparisons, we characterised genetic diversity within each of 16 populations in our cohort defined by country of origin and species by computing a variety of summary statistics using SNP data from the whole genome. These statistics included nucleotide diversity (*θ_π_*; Figure 5a), the density of segregating sites (*θ_W_*; Figure S5), Tajima’s *D* (Figure 5b) and site frequency spectra (SFS; Figure S6). We also estimated runs of homozygosity (ROH; Figure 5c) within each individual and runs of identity by descent (IBD; Figure 5d) between individuals, both of which provide additional information about haplotype sharing and patterns of relatedness within populations.

**Figure 5.**
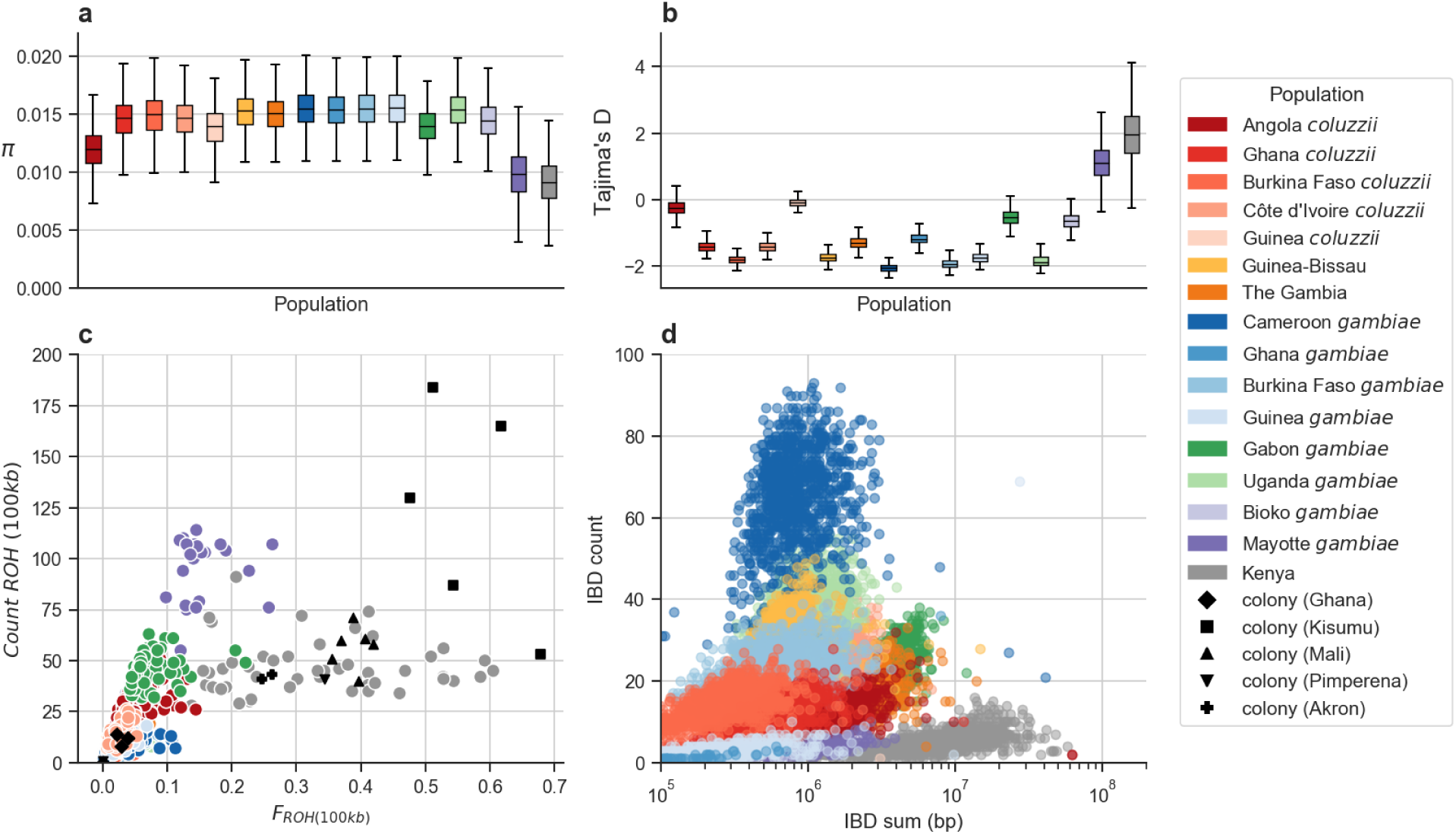
Genetic diversity within populations. **(a)** Nucleotide diversity (*θ_π_*) calculated in non-overlapping 20 kbp genomic windows using SNPs from euchromatic regions of Chromosome 3. **(b)** Tajima’s *D* calculated in non-overlapping 20 kbp genomic windows using SNPs from euchromatic regions of Chromosome 3. **(c)** Runs of homozygosity (ROH) in individual mosquitoes. Each marker represents an individual mosquito. **(d)** Runs of identity by descent between individuals. Each marker represents a pair of individuals drawn from the same population.

The two easternmost populations (Kenya, Mayotte) were outliers in all statistics calculated, with lower diversity, a deficit of rare variants relative to neutral expectation, and a higher degree of haplotype sharing within and between individuals. The Kenyan population was represented in Ag1000G phase 1, and we previously described how the patterns of diversity in this population were consistent with a severe and recent population bottleneck [12]. The similarities between Kenya and Mayotte suggest that the Mayotte population has also experienced a population bottleneck, which would be expected given that Mayotte is an oceanic island 310 km from Madagascar and 500 km from continental Africa, and may have been colonised by *An. gambiae* via small numbers of individuals. Although ROH and IBD were elevated in both populations, Mayotte individuals had a larger number of shorter tracts than Kenyan individuals, which may reflect differences in the timing and/or strength of a bottleneck. In contrast, the *An. gambiae* individuals from Bioko Island had similar patterns of diversity to *An. gambiae* populations from West and Central Africa, supporting other analyses which suggest that this population is not strongly isolated from continental populations (Figures S2, 3). The additional *An. coluzzii* populations (Ghana, Côte d’Ivoire) were similar to the previously sampled Burkina Faso *An. coluzzii* population, and the newly sampled Gambian population with uncertain species status was similar to the previously sampled Guinea-Bissau population, consistent with evidence from population structure analyses that these populations form groupings with shared demographic histories and ongoing gene flow.

### Design of Cas9 gene drives

Nucleotide variation data from this resource is being used to inform the development of gene drives, a novel mosquito control technology using engineered selfish genetic elements to cause mosquito population suppression or modification [35, 36, 37, 38, 8]. Promising results have been obtained with a Cas9 homing endonuclease gene drive targeting a locus in the doublesex gene (*dsx*), which is a critical component of the sex determination pathway [8]. This locus was chosen in part because it has extremely low genetic diversity both within and between species in the *An. gambiae* complex [12]. Low diversity is required because any natural variation within the target sequence could inhibit association with the Cas9 guide RNA and cause resistance to the gene drive [39]. We reviewed nucleotide variation within *dsx* using the expanded cohort of wild-caught samples in the phase 2 cohort, and found no new nucleotide variants within the sequence targeted for Cas9 gene drive, other than the previously known SNP at 2R:48,714,641, which has been shown not to interfere with the gene drive process in lab populations [8]. To facilitate the search for other potential gene drive targets in *dsx* and other genes, we computed allele frequencies for all SNPs in all populations and included those data in the resource. We also compiled a table of all potential Cas9 target sites (23 bp regions with a protospacer-adjacent motif) in the genome that overlap a gene exon. This table includes a total of 20 Cas9 targets that overlap *dsx* exon 5 and that contain at most one SNP within the Ag1000G phase 2 cohort (Figure 6). Thus there may be multiple viable targets for gene drives disrupting the sex determination pathway, providing opportunities to mitigate the impact of resistance due to variation within any single target.

**Figure 6.**
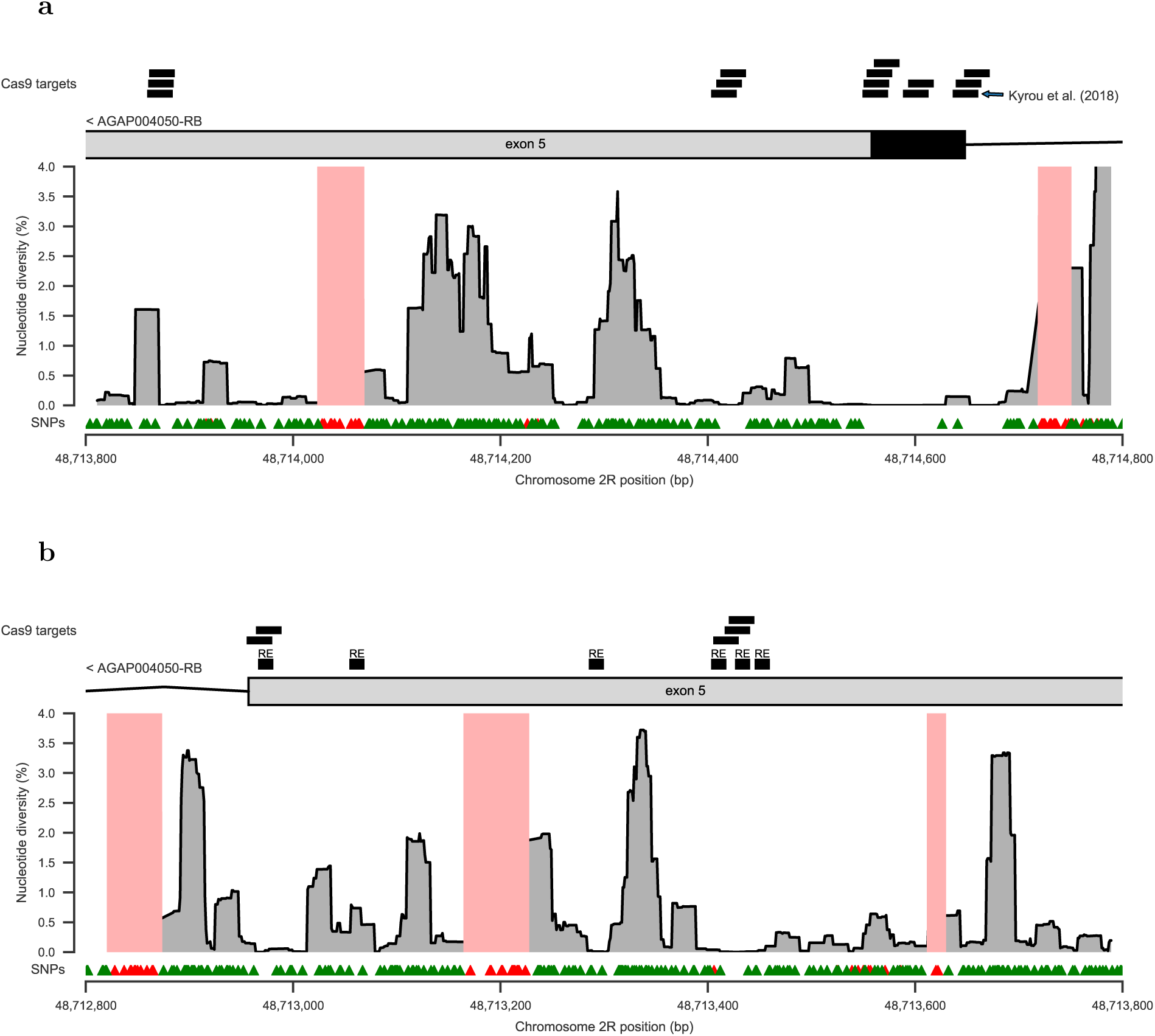
Nucleotide diversity within the female-specific exon 5 of the doublesex gene (*dsx*; AGAP004050), a key component of the sex determination pathway and a gene targeted for Cas9-based homing endonuclease gene drive [8]. In both plots, the location of exon 5 within the female-specific isoform (AGAP004050-RB; AgamP4.12 gene set) is shown above (black = coding sequence; grey = untranslated region), with additional annotations above to show the location of viable Cas9 target sequences containing at most 1 SNP, and the putative exon splice enhancing sequences (“RE”) reported in [40]. The main region of the plot shows nucleotide diversity averaged across all Ag1000G phase 2 populations, computed in 23 bp moving windows. Regions shaded pale red indicate regions not accessible to SNP calling. Triangle markers below show the locations of SNPs discovered in Ag1000G phase 2 (green = passed variant filters; red = failed variant filters). **(a)** exon5/intron4 boundary. **(b)** exon5/intron6 boundary.

The presence of highly conserved regions within *dsx* also provides an example of how genetic variation data from natural populations can be relevant to the study of fundamental molecular processes such as sex determination. The region of conservation containing the Cas9 target site in fact extends over 200 bp, including 50 bp of untranslated sequence within exon 5, the entire coding sequence of exon 5, and 50 bp of intron 4 (Figure 6a). Such conservation of both coding and non-coding sites suggests that purifying selection is acting here on the nucleotide sequence and not just on the protein sequence. This in turn suggests that the nucleotide sequence serves as an important target for factors that bind to DNA or pre-mRNA molecules. This is plausible because sex determination in insects depends on sex-specific splicing of *dsx*, with exon 5 being included in the female transcript and excluded in the male transcript [41]. The upstream regulatory factors that control this differential splicing are not known in *An. gambiae* [40, 42], but in *Drosophila melanogaster* it has been shown that female-specific factors bind to regulatory sequences (*dsxRE* s) within the exon 5 region of the *dsx* pre-mRNA and promote inclusion of exon 5 within the final transcript [43, 41]. Putative homologs of these (*dsxRE*) sequences are present in *An. gambiae* [40], and five out of six *dsxRE* s are located in tracts of near-complete nucleotide conservation in our data, consistent with purifying selection due to pre-mRNA binding (Figure 6b). However, the 200 bp region of conservation spanning the intron 4/exon 5 boundary targeted for Cas9 gene drive remains mysterious, because it is more than 1 kbp distant from any of these putative regulatory sequences. Overall these data add further evidence for fundamental differences in the molecular biology of sex determination between *Anopheles* and *Drosophila* and provide new clues for further investigation of the molecular pathway upstream of *dsx* in *An. gambiae* [40, 42].

### Resistance to pyrethroid insecticides

Malaria control in Africa depends heavily on mass distribution of long-lasting insecticidal bed-nets (LLINs) impregnated with pyrethroid insecticides [44, 45, 46]. Entomological surveillance programs regularly test malaria vector populations for pyrethroid resistance using standardised bioassays, and these data have shown that pyrethroid resistance has become widespread in *An. gambiae* [2, 3]. However, pyrethroid resistance can be conferred by different molecular mechanisms, and it is not well understood which molecular mechanisms are responsible for resistance in which mosquito populations. The nucleotide variation data in this resource include 66 non-synonymous SNPs within the *Vgsc* gene that encodes the binding target for pyrethroid insecticides, of which two SNPs (`L995F, L995S`) are known to confer a pyrethroid resistance phenotype, and one SNP (`N1570Y`) has been shown to substantially increase pyrethroid resistance when present in combination with `L995F` [47]. These SNPs can serve as markers of target-site resistance to pyrethroids, but knowledge of genetic markers of metabolic resistance in *An. gambiae* and *An. coluzzii* is currently limited [48, 49]. Metabolic resistance to pyrethroids is mediated at least in part by increased expression of cytochrome P450 (CYP) enzymes [50, 51, 52, 53], and we found CNV hot-spots at two loci containing *Cyp* genes [19]. One of these loci occurs on chromosome arm 2R and overlaps a cluster of 10 *Cyp* genes, including *Cyp6p3* previously shown to metabolise pyrethroids [54] and recently shown to confer pyrethroid resistance when expression is increased in *An. gambiae* using the GAL4/UAS transgenic system [55]. The second locus occurs on the X chromosome and spans a single *Cyp* gene, *Cyp9k1*, which has also been shown to metabolise pyrethroids [53]. At each of these two loci we found a remarkable allelic heterogeneity, with at least 15 distinct CNV alleles, several of which were present in over 50% of individuals in some populations and were associated with signatures of positive selection [19]. We also found CNVs at two other *Cyp* gene loci on chromosome arm 3R containing genes previously associated with pyrethroid resistance, *Cyp6z1* [56] and *Cyp6m2* [57], although there was only a single CNV allele at each locus. Overexpression of *Cyp6m2* has been shown to confer resistance to pyrethroids but increased susceptibility to the organophosphate malathion [55], and so the selection pressures at this locus may be more complex. The precise phenotype of these CNVs remains to be characterised, but given the multiple lines of evidence showing that increased expression of genes at these loci confers pyrethroid resistance, it seems reasonable to assume that CNVs at these loci can serve as a molecular marker of CYP-mediated metabolic resistance to pyrethroids.

We constructed an overview of the prevalence of these two pyrethroid resistance mechanisms – target-site resistance and CYP-mediated metabolic resistance – within the Ag1000G phase 2 cohort by combining the data on nucleotide and copy number variation (Figure 7). The sampling of these populations was conducted at different times in different locations, and the geographical sampling is relatively sparse, so we cannot draw any general conclusions about the current distribution of resistance from our data. However, some patterns were clear. For example, West African populations of both species (Burkina Faso, Guinea, Côte d’Ivoire) all had more than 84% of individuals carrying both target-site and metabolic resistance markers. In Ghana, Cameroon, Gabon and Angola, target-site resistance was nearly fixed in all populations, but metabolic resistance markers were at lower frequencies, and the samples from Bioko Island carried no resistance markers at all. The Bioko samples were collected in 2002, and so the lack of resistance is likely due to the fact that sampling predated any major scale-up of vector control interventions [53]. However, the Gabon samples were collected in 2000, and show that high levels of target-site resistance were present in some populations at that time. In the “Far West” (Guinea Bissau, The Gambia) [58], target-site resistance was absent, but *Cyp* gene amplifications were present, and thus surveillance using only molecular assays that detect target site resistance at those locations could be missing an important signal of metabolic resistance. In East Africa, both Kenya and Uganda had high frequencies of target-site resistance (88% and 100% respectively). However, 81% of Uganda individuals also had *Cyp* gene amplifications, whereas only 4% of Kenyans (two individuals) carried these metabolic resistance markers. Denser spatio-temporal sampling and sequencing will enable us to build a more complete picture of the prevalence and spread of these different resistance mechanisms, and would be highly relevant to the design of insecticide resistance management plans.

**Figure 7.**
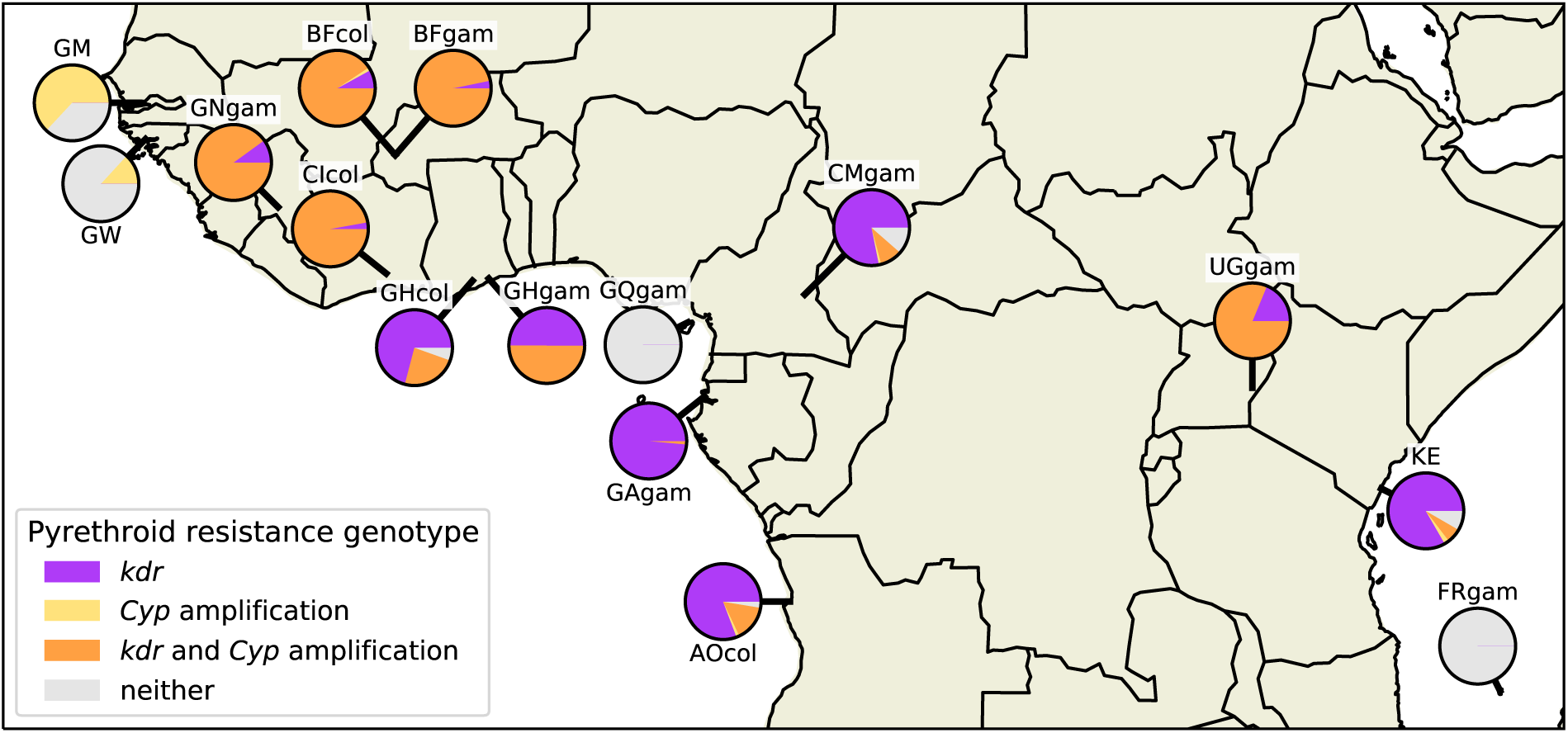
Pyrethroid resistance genotype frequencies. The geographical distribution of pyrethroid insecticide resistance genotypes are shown by population. Pie chart colours represent resistance genotype frequencies: purple - these individuals were either homozygous or heterozygous for one of the two *kdr* pyrethroid target site resistance alleles *Vgsc*-L995F/S; yellow - these individuals carried a copy number amplification within any of the *Cyp6p/aa*, *Cyp6m*, *Cyp6z* or *Cyp9k* gene clusters, but no *kdr* alleles; orange - these individuals carried at least one *kdr* allele and one *Cyp* gene amplification; grey - these individuals carried no known pyrethroid resistance alleles (no *kdr* alleles or *Cyp* amplifications). The Guinea *An. coluzzii* population is omitted due to small sample size.

## Discussion

### Insecticide resistance surveillance

The Ag1000G phase 2 data resource incorporates both nucleotide and copy number variation from the whole genomes of 1,142 mosquitoes collected from 13 countries spanning the African continent. These data provide a battery of new genetic markers that can be used to expand our capabilities for molecular surveillance of insecticide resistance. Insecticide resistance management is a major challenge for malaria vector control, but the availability of new vector control products is opening up new possibilities. However, new products may be more expensive than products currently in use, so procurement decisions have to be justified, and resources targeted to areas where they will have the greatest impact. For example, next-generation LLINs are now available which combine a pyrethroid insecticide with either a second insecticide or a synergist compound, piperonyl butoxide (PBO), which partially ameliorates metabolic resistance by inhibiting CYP enzyme activity in the mosquito. However, CYP-mediated metabolic resistance is only one of several possible mechanisms of pyrethroid resistance that may or may not be present in vector populations being targeted. It would therefore be valuable to survey mosquito populations and determine the prevalence of different pyrethroid resistance mechanisms, both before and after any change in vector control strategy. Our data resource includes CNVs at four *Cyp* gene loci (*Cyp6p/aa*, *Cyp6m*, *Cyp6z* and *Cyp9k*) which could serve as molecular markers of CYP-mediated metabolic resistance. Glutathione S-transferase enzymes are also associated with metabolic resistance to pyrethroids [59, 55] as well as to other insecticide classes [48, 60, 61, 55] and we found CNVs at the *Gste* locus which could serve as molecular markers of this alternative resistance mechanism, which is not inhibited by PBO. *Gste* CNVs were less prevalent in our dataset than *Cyp* CNVs, and the geographical distribution also differed, suggesting they may be driven by different selection pressures (Figure S7). Further work is needed to characterise the resistance phenotype associated with these CNVs, but the allelic heterogeneity, the high population frequencies, and the evidence for positive selection observed in our data, coupled with previous gene expression and functional studies [50, 51, 52, 53, 55], all support a metabolic role in insecticide resistance.

To illustrate the potential for improved molecular surveillance of pyrethroid resistance, we combined the data on known SNP markers of target-site resistance and the novel putative CNV markers of CYP-mediated metabolic resistance, and computed the frequencies of these different resistance mechanisms in the populations we sampled (Figure 7). There are clear heterogeneities, with some populations at high frequency for both resistance mechanisms, particularly in West Africa. The presence of CYP-mediated pyrethroid resistance in a population suggests that PBO LLINs might provide some benefit over standard LLINs. However, if other resistance mechanisms are also at high frequency, the benefit of the PBO synergist might be diminished. Current WHO guidance states that PBO LLINs are recommended in regions with “intermediate levels” of pyrethroid resistance, but not where resistance levels are high [62]. This guidance is based on modelling of bioassay data and experimental hut trials, and it is not clear why PBO LLINs are predicted to provide diminishing returns at higher resistance levels, although high levels of resistance presumably correlate with the presence of multiple resistance mechanisms, including mechanisms not inhibited by PBO [45]. Without molecular data, however, this guidance is hard to evaluate or improve upon.

Ideally, molecular data on insecticide resistance mechanisms would be collected as part of routine entomological surveillance, as well as in field trials of new vector control products, alongside data from bioassays and other standard entomological monitoring procedures. There are several options for scaling up surveillance of new genetic markers, including both whole genome sequencing and targeted (amplicon) sequencing with several choices of sequencing technology platform, as well as various PCR-based assays. Assays that target specific genetic loci are attractive in the short term, because of the low cost and infrastructure requirements, and data from the Ag1000G project have been used successfully to design multiplex assays for the Agena Biosciences iPLEX platform [63] and for Illumina amplicon sequencing (manuscript in preparation). But targeted assays would need to be updated regularly to ensure all current forms of insecticide resistance are covered, and to capture new forms of resistance as they emerge. None of the samples sequenced in this study were collected more recently than 2012, geographical sampling within each country was limited, and many countries are not yet represented in the resource, therefore there remain important gaps to be filled. The next phase of the Ag1000G project will expand the resource to cover 18 countries, and will include another major malaria vector, *An. arabiensis*, in addition to *An. gambiae* and *An. coluzzii*, and so will address some of these gaps. Looking beyond the Ag1000G project, genomic surveilance of insecticide resistance will require new sampling frameworks that incorporate spatial and ecological modelling of vector distributions to improve future collections and guide sampling at appropriate spatial scales [64]. To keep pace with vector populations, regular whole genome sequencing of contemporary populations from a well-chosen set of sentinel sites will be needed. Fortunately mosquitoes are easy to transport, and the costs of whole genome sequencing continue to fall, so it is reasonable to consider a mixed strategy that includes both whole genome sequencing and targeted assays.

### Gene flow

These data also cast some new, and in some cases contrasting, light on the question of gene flow between malaria vector populations. The question is of practical interest because gene flow is enabling the spread of insecticide resistance between species and across large geographical distances [12, 65]. This gene flow also needs to be quantified and modelled before new vector control interventions based on the release of genetically modified mosquitoes could be considered [66]. We found evidence that isolation by distance is greater for *An. coluzzii* than for *An. gambiae*, at least within West Africa, suggesting that the effective rate of migration could be lower in *An. coluzzii*. This result was supported by population structure analyses, where all *An. coluzzii* individuals were clearly clustered by country in the UMAP analysis, whereas *An. gambiae* individuals from Guinea, Burkina Faso, Ghana and Cameroon could not be separated in any of the UMAP, PCA or admixture analyses. A variety of anopheline species have recently been found to engage in long-distance wind-assisted migration, including *An. coluzzii* but not *An. gambiae*, which would appear to contradict our results, although the study was limited to a single location within the Sahelian region [32]. If *An. coluzzii* does have a lower rate and/or range of dispersal than *An. gambiae*, this is clearly not limiting the spread of insecticide resistance adaptations between countries. For example, among the CNV alleles we discovered at the *Cyp6p/aa*, *Cyp9k1* and *Gste* loci, 7/13 alleles found in *An. coluzzii* had spread to more than one country, compared with 8/27 alleles in *An. gambiae* [19]. There is also an interesting contrast between the spread of pyrethroid target-site and metabolic resistance alleles. Our previous analysis of haplotypes carrying target-site resistance alleles in the Ag1000G phase 1 cohort found that resistance haplotypes had spread to countries spanning the equatorial rainforest and the Rift valley, and had moved between *An. gambiae* and *An. coluzzii* [12, 65]. In the most extreme example, one haplotype (F1) had spread to countries as distant as Guinea and Angola. In contrast, although CNV alleles were commonly found in multiple countries, we did not observe any cases of CNV alleles crossing any of these ecological or biological boundaries, apart from a single allele found in both Gabon and Cameroon *An. gambiae* (*Gste* Dup5). There are multiple possible explanations for this difference, including differences in the strength, timing or spatial distribution of selective pressures, or intrinsic factors such as differences in fitness costs in the absence of positive selection. Further work is required to investigate the selective forces and biological constraints affecting the spread of these different modes of adaptation to insecticide use. The two island populations sampled in this project phase also provide an interesting contrast. Samples from Mayotte are highly differentiated from mainland *An. gambiae* and have patterns of reduced genetic diversity, consistent with a reduction in population size and strong isolation. Bioko samples, on the other hand, are closely related to West African *An. gambiae*, and have comparable levels of genetic diversity, suggesting ongoing gene flow. Bioko is part of Equatorial Guinea administratively, and there are frequent ferries to the mainland, which could provide opportunities for mosquito movement. However, there are no pyrethroid resistance alleles in our Bioko samples and these were collected in 2002 at a time when target-site resistance alleles were present in mainland populations, so the rate of contemporary migration between Bioko and mainland populations remains an open question. A recent study of *An. gambiae* populations on the Lake Victoria islands, separated from mainland Uganda by 4-50 km, found evidence for isolation between island and mainland populations, as well as between individual islands [67]. However, some selective sweeps at insecticide resistance loci had spread through both mainland and island populations, thus isolation is not complete and some contemporary gene flow occurs. Resolving these gene flow questions and apparent contradictions will require fitting quantitative models of contemporary migration to genomic data. We previously fitted migration models to pairs of populations using site frequency spectra, but the approach provides poor resolution to differentiate recent from ancient migration rates [12]. In general, methods that leverage information about haplotype sharing within and between populations should provide the greatest resolution to disentangle ancient from recent demographic events, as well as providing independent estimates for both migration rates and population densities. There is promising recent work in this direction [68], but models have so far only been applied to data from human populations. The haplotype data we have generated should prove a useful resource for further work to evaluate whether these models can be applied to malaria vector populations with sufficient accuracy to support real-world planning of new vector control interventions.

### Conclusions

Malaria has become a stubborn foe, frustrating global efforts towards elimination in both low and high burden settings. However, new vector control tools offer hope, as does the renewed focus on improving surveillance systems and using data to tailor interventions. The genomic data resource we have generated provides a platform from which to accelerate these efforts, demonstrating the potential for data integration on a continental scale. Nevertheless, work remains to fill gaps in these data, by expanding geographical coverage, including other malaria vector species and integrating genomic data collection with routine surveillance of contemporary populations using quantitative sampling design. We hope that the MalariaGEN data-sharing community and framework for international collaboration can continue to serve as a model for coordinated action.

## Methods

### Population sampling

Ag1000G phase 2 mosquitoes were collected from natural populations at 33 sites in 13 sub-Saharan African countries (Figure 1 & Table S1). Throughout, we use species nomenclature following Coetzee *et al*. [13]; prior to Coetzee *et al*., *An. gambiae* was known as *An. gambiae sensu stricto* (S form) and *An. coluzzii* was known as *An. gambiae sensu stricto* (M form). Details of the eighteen collection sites novel to Ag1000G phase 2 (dates, collection and DNA extraction methods) can be found below. Information pertaining to the collection of samples released as part of Ag1000G phase 1 can be found in the supplementary information of [12]. Unless otherwise stated, the DNA extraction method used for the collections described below was Qiagen DNeasy Blood and Tissue Kit (Qiagen Science, MD, USA).

#### Côte d’Ivoire

Tiassalé (5.898, −4.823) is located in the evergreen forest zone of southern Côte d’Ivoire. The primary agricultural activity is rice cultivation in irrigated fields. High malaria transmission occurs during the rainy seasons, between May and November. Samples were collected as larvae from irrigated rice fields by dipping between May and September 2012. All larvae were reared to adults and females preserved over silica for DNA extraction. Specimens from this site were all *An. coluzzii*, determined by PCR assay [21].

#### Bioko

Collections were performed during the rainy season in September, 2002 by overnight CDC light traps in Sacriba of Bioko island (3.7, 8.7). Specimens were stored dry on silica gel before DNA extraction. Specimens contributed from this site were *An. gambiae* females, genotype determined by two assays [69, 70]. All specimens had the 2L^+a^/2L^+a^ karyotype as determined by the molecular PCR diagnostics [71]. These mosquitoes represent a population that inhabited Bioko Island before a comprehensive malaria control intervention initiated in February 2004 [72]. After the intervention *An. gambiae* was declining, and more recently almost only *An. coluzzii* can be found [73].

#### Mayotte

Samples were collected as larvae during March-April 2011 in temporary pools by dipping, in Bouyouni (−12.738, 45.143), M’Tsamboro Forest Reserve (−12.703, 45.081), Combani (−12.779, 45.143), Mtsanga Charifou (−12.991, 45.156), Karihani Lake forest reserve (−12.797, 45.122), Mont Benara (−12.857, 45.155) and Sada (−12.852, 45.104) in Mayotte island. Larvae were stored in 80% ethanol prior to DNA extraction. All specimens contributed to Ag1000G phase 2 were *An. gambiae* [70] with the standard 2L^+a^/2L^+a^ or inverted 2L^a^/2L^a^ karyotype as determined by the molecular PCR diagnostics [71]. The samples were identified as males or females by the sequencing read coverage of the X chromosome using LookSeq [74].

#### The Gambia

Indoor resting female mosquitoes were collected by pyrethrum spray catch from four hamlets around Njabakunda (−15.90, 13.55), North Bank Region, The Gambia between August and October 2011. The four hamlets were Maria Samba Nyado, Sare Illo Buya, Kerr Birom Kardo, and Kerr Sama Kuma; all are within 1 km of each other. This is an area of unusually high rates of apparent hybridization between *An. gambiae s.s.* and *An. coluzzii* [75, 76]. Njabakunda village is approximately 30 km to the west of Farafenni town and 4 km away from the Gambia River. The vegetation is a mix of open savannah woodland and farmland.

#### Ghana

Mosquitoes were collected from Twifo Praso (5.609, −1.549), a peri-urban community located in semi-deciduous forest in the Central Region of Ghana. It is an extensive agricultural area characterised by small-scale vegetable growing and large-scale commercial farms such as oil palm and cocoa plantations. Mosquito samples were collected as larvae from puddles near farms between September and October, 2012. Madina (5.668, −0.219) is a suburb of Accra within the coastal savanna zone of Ghana. It is an urban community characterised by numerous vegetable-growing areas. The vegetation consists of mainly grassland interspersed with dense short thickets often less than 5 m high with a few trees. Specimens were sampled from puddles near roadsides and farms between October and December 2012. Takoradi (4.912, −1.774) is the capital city of Western Region of Ghana. It is an urban community located in the coastal savanna zone. Mosquito samples were collected from puddles near road construction and farms between August and September 2012. Koforidua (6.094, −0.261) is the capital city of Eastern Region of Ghana and is located in semi-deciduous forest. It is an urban community characterized by numerous small-scale vegetable farms. Samples were collected from puddles near road construction and farms between August and September 2012. Larvae from all collection sites were reared to adults and females preserved over silica for DNA extraction. Both *An. gambiae* and *An. coluzzii* were collected from these sites, determined by PCR assay [21].

#### Guinea-Bissau

Mosquitoes were collected in October 2010 using indoor CDC light traps, in the village of Safim (11.957, −15.649), ca. 11 km north of Bissau city, the capital of the country. Malaria is hyperendemic in the region and transmitted by members of the *Anopheles gambiae* complex [77]. *An. arabiensis*, *An. melas*, *An. coluzzii* and *An. gambiae*, as well as apparent hybrids between the latter two species, are known to occur in the region [78, 77]. Mosquitoes were preserved individually on 0.5ml micro-tubes filled with silica gel and cotton. DNA extraction was performed by a phenol-chloroform protocol [79].

### Lab crosses

The Ag1000G phase 2 data release includes the genomes of seven additional lab colony crosses, both parents and offspring (Table S2): cross 18-5 (Ghana mother x Kisumu/G3 father, 20 offspring); 37-3 (Kisumu x Pimperena, 20 offspring); 45-1 (Mali x Kisumu, 20 offspring); 47-6 (Mali x Kisumu, 20 offspring); 73-2 (Akron x Ghana, 19 offspring); 78-2 (Mali x Kisumu/Ghana, 19 offspring); 80-2 (Kisumu x Akron, 20 offspring). Father colonies with two names, e.g., “Kisumu/G3”, signify that the father is from one of these two colonies, but exactly which one is unknown. The colony labels, e.g., “18-5”, are identifiers used for each of the crosses within the project and have no particular meaning. Information pertaining to the crosses released as part of Ag1000G phase 1 can be found in the supplementary information of [12] as well as methods for cross creation and processing that also apply to the crosses in phase 2.

### Whole genome sequencing

Sequencing was performed on the Illumina HiSeq 2000 platform at the Wellcome Sanger Institute. Paired-end multiplex libraries were prepared using the manufacturer’s protocol, with the exception that genomic DNA was fragmented using Covaris Adaptive Focused Acoustics rather than nebulization. Multiplexes comprised 12 tagged individual mosquitoes and three lanes of sequencing were generated for each multiplex to even out variations in yield between sequencing runs. Cluster generation and sequencing were undertaken per the manufacturer’s protocol for paired-end 100 bp sequence reads with insert size in the range 100-200 bp. Target coverage was 30X per individual.

### Genome accessibility

For various population-genomic analyses, it is necessary to have a map of which positions in the reference genome can be considered accessible, at which we can confidently call nucleotide variation. For Ag1000G phase 2, we repeated the phase 1 genome acccessibility analyses [12] with 1,142 samples and the additional Mendelian error information provided by the 11 crosses (in phase 1 there were four crosses). These analyses constructed a number of annotations for each position in the reference genome, based on data from sequence read alignments from all wild-caught samples, and additional data from repeat annotations. These annotations were then analysed for their association with rates of variants with one or more Mendelian errors in the crosses. Annotations and thresholds were chosen to remove classes of variants that were enriched for Mendelian errors. Following these analyses it was apparent that the accessibility classifications used in Ag1000G phase 1 were also appropriate in application to phase 2. Reference genome positions were classificed as accessible if: Not repeat masked by DUST; No Coverage <= 0.1% (at most 1 individual had zero coverage); Ambiguous Alignment <= 0.1% (at most 1 individual had ambiguous alignments); High Coverage <= 2% (at most 20 individuals had more than twice their genome-wide average coverage); Low Coverage <= 10% (at most 114 individuals had less than half their genome-wide average coverage); Low Mapping Quality <= 10% (at most 114 individuals had average mapping quality below 30).

We performed additional analyses to verify that there was no significant bias towards one species or another given the use of a single reference genome AgamP3 [9] for alignment of reads from all individuals. We found that the genomes of *An. coluzzii* and *An. gambiae* individuals were similarly diverged from the reference genome (Fig. S8). The similarity in levels of divergence is likely to reflect the mixed ancestry of the PEST strain from which the reference genome was derived [9, 10]. An exception to this was the pericentromeric region of the X chromosome, a known region of divergence between the two species [12] where the reference genome is closer to *An. coluzzii* than to *An. gambiae*. The similarity of this region to *An. coluzzii* may be due to artificial selection for the X-linked pink eye mutation in the reference strain [9], as this originated in the *An. coluzzii* parent it may have led to the removal of any *An. gambiae* ancestry in this region.

### Sequence analysis and variant calling

SNP calling methods were unchanged from phase 1 of the Anopheles 1000 genomes project [12]. Briefly, sequence reads were aligned to the AgamP3 reference genome [9, 10] using bwa version 0.6.2, duplicate reads marked [80], reads realigned around putative indels, and SNPs discovered using GATK version 2.7.4 Unified Genotyper following best practice recommendations [81].

### Sample quality control

A total of 1,285 individual mosquitoes were sequenced as part of Ag1000G phase 2 and included in the cohort for variant discovery. After variant discovery, quality-control (QC) steps using coverage and contamination filters alongside principal component analysis and metadata concordance were performed to exclude individuals with poor quality sequence and/or genotype data as detailed in [12]. A total of 143 individuals were excluded at this stage, retaining 1,142 individuals for downstream analyses. Any SNPs with variant alleles found only in excluded samples were then also excluded.

### Variant Filtering

Following Ag1000G phase 1 [12], we applied the following SNP filters to reduce the number of false SNP discoveries. We filtered any SNP that occurred at a genome position classified as inaccessible as described in the section on genome accessibility above, thus removing SNPs with evidence for excessively high or low coverage or ambiguous alignment. We then applied additional filters using variant annotations produced by GATK based on an analysis of Mendelian error in all 11 crosses present in phase 2 and Ti/Tv ratio, similar to that described above for the genome accessibility analysis. We filtered any SNP that failed any of the following criteria: QD <5; FS >100; ReadPosRankSum <-8; BaseQRankSum <-50.

Of 105,486,698 SNPs reported in the raw callset, 57,837,885 passed all quality filters, 13,760,984 (23.8%) of which were multi-allelic (three or more non-reference alleles). To produce an analysis-ready VCF file for each chromosome arm, we first removed all non-SNP variants. We then removed genotype calls for individuals excluded by the sample QC analysis described above, then removed any variants that were no longer variant after excluding individuals. We then added INFO annotations with genome accessibility metrics and added FILTER annotations per the criteria defined above. Finally, we added INFO annotations with information about functional consequences of mutations using SNPEFF version 4.1b [82].

### Haplotype estimation

Haplotype estimation, also known as phasing, was performed on all phase 2 wild-caught individuals using unchanged methodology from phase 1 of the Anopheles 1000 genomes project [12]. In short, SHAPEIT2 was used to perform statistical phasing with information from sequence reads [83].

### Population structure

Ancestry informative marker (AIM), *F_ST_*, doubleton sharing and SNP PCA were conducted following methods defined in [12]. The PCA and UMAP analyses were performed on 131,679 SNPs from euchromatic regions of chromosome arms 3L and 3R obtained from the full dataset via random downsampling to 100,000 non-singleton SNPs from each chromosome arm then performing LD-pruning. To generate the UMAP projection shown in Figure 2, each country and species was downsampled to a maximum of 50 individuals, to provide a projection that was less warped by differences in sample size. The UMAP analysis was also performed on the full set of individuals, which gave qualitatively identical results in terms of the clustering of individuals. UMAP was performed using the umap-learn Python package [26] with the following parameter settings: *n*_*neighbors* = 15; *min*_*dist* = 2; *spread* = 5; *metric* = *euclidean*. Other parameter values for *n*_*neighbours* and *min*_*dist* were also performed, all producing qualitatively identical results. One population (Guinea *An. coluzzii*, n=4) was excluded from *F_ST_* analysis and three populations (Guinea *An. coluzzii*, n=4; Bioko *An. gambiae*, n=9; Ghana *An. gambiae*, n=12) were excluded from doubleton sharing analysis due to small sample size. All analyses of geographical population structure using SNP data were conducted on euchromatic regions of Chromosome 3 (3R:1-37 Mbp, 3L:15-41 Mbp), which avoids regions of polymorphic inversions, reduced recombination and unequal divergence from the reference genome [12]. Unscaled CNV variation PCAs were built from the CNV presence/absence calls [19], using the *prcomp* function in R [84].

Admixture models were fitted using the program LEA version 2.0 [85] in R version 3.6.1 [84]. Ten independent sets of SNPs were generated by selecting SNPs from euchromatic regions of Chromosome 3 with minor allele frequency greater than 1%, then randomly selecting 100,000 SNPs from each chromosome arm, then applying the same LD pruning methodology as used for PCA, then combining back together remaining SNPs from both chromosome arms. The resulting files were exported in .geno format, which were then analyzed using the *snmf* method (sparse non-negative matrix factorization [28]) to obtain ancestry estimates to each cluster (K) tested. We tested all K values from 2 to 15. Ten replicates of the analysis with *snmf* were run for each dataset, which meant that 100 runs were performed for each K. We assessed the convergence and replicability of the results across the 100 runs (ten different datasets, each one replicated ten times dataset) using CLUMPAK [86]. CLUMPAK was used to summarize the results, identify the major and minor clustering solutions identified at each K (if they occurred), and estimate the average ancestry proportions for the major solution which was used to interpret the results. We assessed how the clustering solution fitted with the data using the cross-entropy criterion. The lower this criterion is, the better is the model fit to the data.

### Genetic diversity

Analyses of genetic diversity, including nucleotide diversity, Tajima’s D, ROH and IBD (identity by descent), were conducted following methods defined in [12] but using the phase 2 data release of 1,142 samples. In short, scikit-allel version 1.2.0 was used to calculate windowed averages of nucleotide diversity and Tajima’s D [87], IBDseq version r1206 [88] was used to calculate IBD, and an HMM implemented in scikit-allel was used to calculate ROH.

## The *Anopheles gambiae* 1000 Genomes Consortium

Chris S. Clarkson and Alistair Miles jointly led curation of the phase 2 data resource and wrote the paper.

### Data analysis group

Chris S. Clarkson^1^, Alistair Miles^2,1^, Nicholas J. Harding^2^, Eric R. Lucas^3^, C. J. Battey^4^, Jorge Edouardo Amaya-Romero^5,6^, Andrew D. Kern^4^, Michael C. Fontaine^5,6^, Martin J. Donnelly^3,1^, Mara K. N. Lawniczak^1^ and Dominic P. Kwiatkowski^1,2^ (chair).

### Partner working group

Martin J. Donnelly^3,1^ (chair), Diego Ayala^7,5^, Nora J. Besansky^8^, Austin Burt^9^, Beniamino Caputo^10^, Alessandra della Torre^10^, Michael C. Fontaine^5,6^, H. Charles J. Godfray^11^, Matthew W. Hahn^12^, Andrew D. Kern^4^, Dominic P. Kwiatkowski^2,1^, Mara K. N. Lawniczak^1^, Janet Midega^13^, Samantha O’Loughlin^9^, João Pinto^14^, Michelle M. Riehle^15^, Igor Sharakhov^16,17^, Daniel R. Schrider^18^, Kenneth D. Vernick^19^, David Weetman^3^, Craig S. Wilding^20^ and Bradley J. White^21^.

### Population sampling

**Angola**: Arlete D. Troco^22^, João Pinto^14^; **Bioko**: Jorge Cano^23^; **Burkina Faso**: Abdoulaye Diabaté^24^, Samantha O’Loughlin^9^, Austin Burt^9^; **Cameroon**: Carlo Costantini^5,25^, Kyanne R. Rohatgi^8^, Nora J. Besansky^8^; **Côte d’Ivoire**: Edi Constant^26^, David Weetman^3^; **Gabon**: Nohal Elissa^27^, João Pinto^14^; **Gambia**: Davis C. Nwakanma^28^, Musa Jawara^28^; **Ghana**: John Essandoh^29^, David Weetman^3^; **Guinea**: Boubacar Coulibaly^30^, Michelle M. Riehle^15^, Kenneth D. Vernick^19^; **Guinea-Bissau**: João Pinto^14^, João Dinis^31^; **Kenya**: Janet Midega^13^, Charles Mbogo^13^, Philip Bejon^13^; **Mayotte**: Gilbert Le Goff^5^, Vincent Robert^5^; **Uganda**: Craig S. Wilding^20^, David Weetman^3^, Henry D. Mawejje^32^, Martin J. Donnelly^3^; **Lab crosses**: David Weetman^3^, Craig S. Wilding^20^, Martin J. Donnelly^3^.

### Sequencing and data production

Jim Stalker^33^, Kirk A. Rockett^2^, Eleanor Drury^1^, Daniel Mead^1^, Anna E. Jeffreys^2^, Christina Hubbart^2^, Kate Rowlands^2^, Alison T. Isaacs^3^, Dushyanth Jyothi^34^, Cinzia Malangone^34^ and Maryam Kamali^35,16^.

### Project coordination

Victoria Simpson^2^, Christa Henrichs^2^ and Dominic P. Kwiatkowski^1,2^.

^1^Parasites and Microbes Programme, Wellcome Sanger Institute, Hinxton, Cambridge CB10 1SA, UK.

^2^MRC Centre for Genomics and Global Health, University of Oxford, Oxford OX3 7BN, UK.

^3^Department of Vector Biology, Liverpool School of Tropical Medicine, Pembroke Place, Liverpool L3 5QA, UK.

^4^Institute for Ecology and Evolution, University of Oregon, 301 Pacific Hall, Eugene, OR 97403, USA.

^5^Laboratoire MIVEGEC (Université de Montpellier, CNRS 5290, IRD 229), Centre IRD de Montpellier, 911, Avenue Agropolis BP 64501, 34395 Montpellier Cedex 5, France.

^6^Groningen Institute for Evolutionary Life Sciences (GELIFES), University of Groningen, PO Box 11103 CC, Groningen, The Netherlands.

^7^Unit d’Ecologie des Systèmes Vectoriels, Centre International de Recherches Médicales de Franceville, Franceville, Gabon.

^8^Eck Institute for Global Health, Department of Biological Sciences & University of Notre Dame, IN 46556, USA.

^9^Department of Life Sciences, Imperial College, Silwood Park, Ascot, Berkshire SL5 7PY, UK.

^10^Istituto Pasteur Italia âĂŞ Fondazione Cenci Bolognetti, Dipartimento di Sanita Pubblica e Malattie Infettive, Università di Roma SAPIENZA, Rome, Italy.

^11^Department of Zoology, University of Oxford, 11a Mansfield Road, Oxford OX1 3SZ, UK.

^12^Department of Biology and School of Informatics and Computing, Indiana University, Bloomington, IN 47405, USA.

^13^KEMRI-Wellcome Trust Research Programme, PO Box 230, Bofa Road, Kilifi, Kenya.

^14^Global Health and Tropical Medicine, GHTM, Instituto de Higiene e Medicina Tropical, IHMT, Universidade Nova de Lisboa, UNL, Rua da Junqueira 100, 1349-008 Lisbon, Portugal.

^15^Department of Microbiology and Immunology, Medical College of Wisconsin, Milwaukee, WI 53226, USA.

^16^Department of Entomology, Virginia Tech, Blacksburg, VA 24061, USA.

^17^Department of Cytology and Genetics, Tomsk State University, Tomsk 634050, Russia.

^18^Department of Genetics, University of North Carolina, 5111 Genetic Medicine Building, 7264, Chapel Hill, NC 27599-7264, USA.

^19^Unit for Genetics and Genomics of Insect Vectors, Institut Pasteur, Paris, France.

^20^School of Biological and Environmental Sciences, Liverpool John Moores University, Liverpool L3 3AF, UK.

^21^Verily Life Sciences, 269 E Grand Ave, South San Francisco, CA 94080, USA.

^22^Programa Nacional de Controle da Malária, Direcção Nacional de Saúde Pública, Ministério da Saúde, Luanda, Angola.

^23^London School of Hygiene & Tropical Medicine. Keppel St, Bloomsbury, London WC1E 7HT, UK.

^24^Institut de Recherche en Sciences de la Santé (IRSS), Bobo Dioulasso, Burkina Faso.

^25^Laboratoire de Recherche sur le Paludisme, Organisation de Coordination pour la lutte contre les Endémies en Afrique Centrale (OCEAC), Yaoundé, Cameroon.

^26^Centre Suisse de Recherches Scientifiques. Yopougon, Abidjan - 01 BP 1303 Abidjan, Côte d’Ivoire.

^27^Institut Pasteur de Madagascar, Avaradoha, BP 1274, 101, Antananarivo, Madagascar.

^28^Medical Research Council Unit The Gambia at the London School of Hygiene & Tropical Medicine (MRCG at LSHTM), Atlantic Boulevard, Fajara, P.O. Box 273, Banjul, The Gambia.

^29^Department of Wildlife and Entomology, University of Cape Coast, Cape Coast, Ghana. ^30^Malaria Research and Training Centre, Faculty of Medicine and Dentistry, University of Mali. ^31^Instituto Nacional de Saaúde Paública, Ministaério da Saaúde Paública, Bissau, Guinaé-Bissau.

^32^Infectious Diseases Research Collaboration, 2C Nakasero Hill Road, PO Box 7475, Kampala, Uganda.

^33^Microbiotica Limited, Biodata, Innovation Centre, Wellcome Genome Campus, Cambridge, CB10 1DR, UK.

^34^European Bioinformatics Institute, Hinxton, Cambridge CB10 1SA, UK.

^35^Department of Medical Entomology and Parasitology, Faculty of Medical Sciences, Tarbiat Modares University, Tehran, Iran.

## Acknowledgments

The authors would like to thank the staff of the Wellcome Sanger Institute Sample Logistics, Sequencing and Informatics facilities for their contributions. The sequencing, analysis, informatics and management of the *Anopheles gambiae* 1000 Genomes Project are supported by Wellcome through Sanger Institute core funding (098051), core funding to the Wellcome Centre for Human Genetics (203141/Z/16/Z), and a strategic award (090770/Z/09/Z); and by the MRC Centre for Genomics and Global Health which is jointly funded by the Medical Research Council and the Department for International Development (DFID) (G0600718; M006212). M.K.N.L. was supported by MRC grant G1100339. S.O.’L. and A.B. were supported by a grant from the Foundation for the National Institutes of Health through the Vector-Based Control of Transmission: Discovery Research (VCTR) program of the Grand Challenges in Global Health initiative of the Bill and Melinda Gates Foundation. D.W., C.S.W., H.D.M. and M.J.D. were supported by Award Numbers U19AI089674 and R01AI082734 from the National Institute of Allergy and Infectious Diseases (NIAID). The content is solely the responsibility of the authors and does not necessarily represent the official views of the NIAID or NIH.

## Data availability

Sequence read alignments and variant calls from Ag1000G phase 2 are available from the European Nucleotide Archive under study accession PRJEB36277 (ENA - http://www.ebi.ac.uk/ena). Sequence read alignments for samples in Ag1000G phase 1 are available under study accession PRJEB18691.

All variation data from Ag1000G phase 2 can also be downloaded from the Ag1000G public FTP site via the MalariaGEN website (https://www.malariagen.net/resource/27).

## Supplementary figures and tables

**Figure S1.**
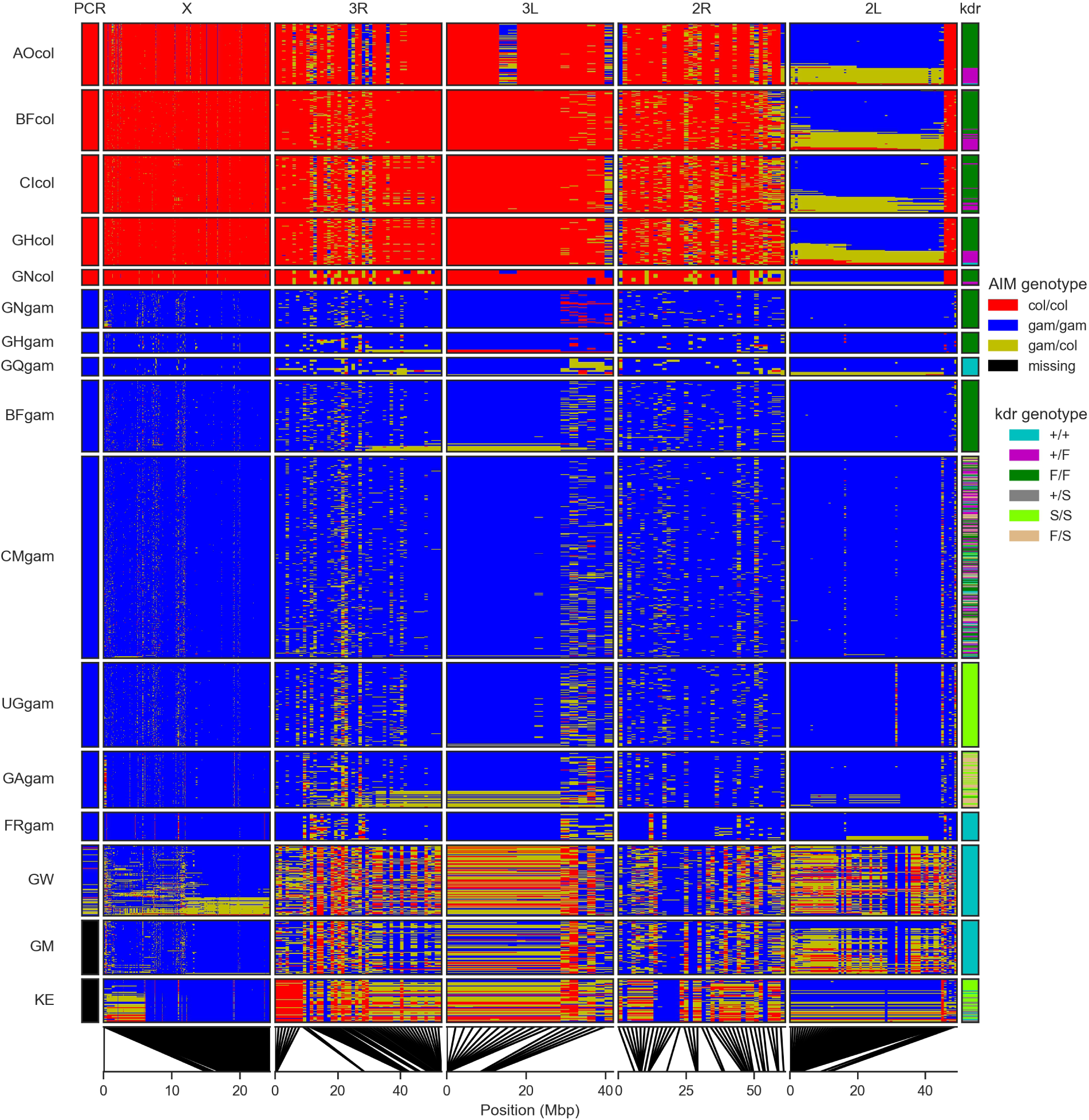
Ancestry informative markers (AIM). Rows represent individual mosquitoes (grouped by population) and columns represent SNPs (grouped by chromosome arm). Colours represent species genotype. The column at the far left (“PCR”) shows the species assignment according to the conventional molecular test based on a single marker on the X chromosome, which was performed for all populations except The Gambia (GM) and Kenya (KE). The column at the far right shows the genotype for *kdr* variants in *Vgsc* codon 995. Lines at the lower edge show the physical locations of the AIM SNPs.

**Figure S2.**
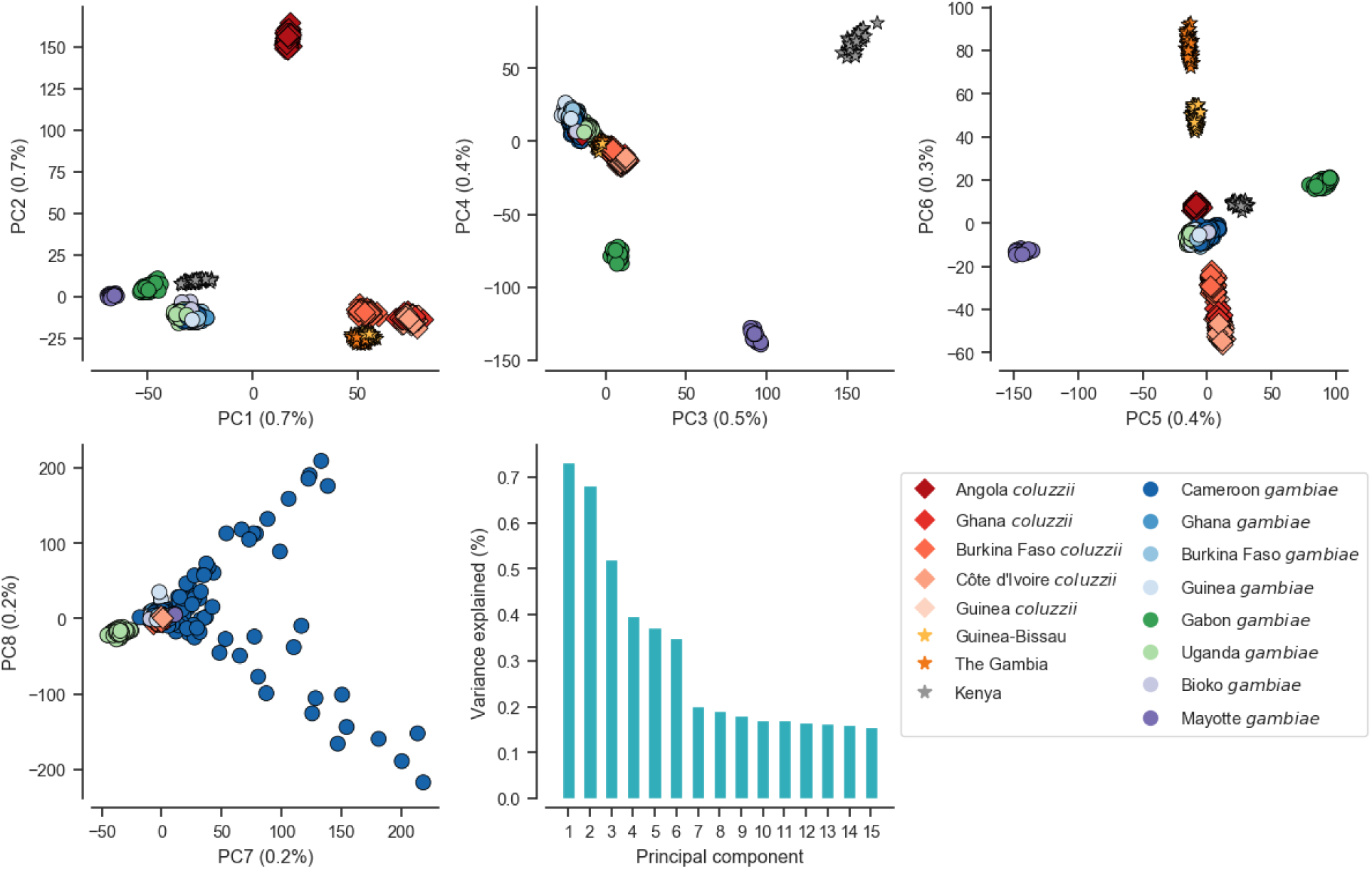
Principal component analysis of the 1,142 wild-caught mosquitoes using biallelic SNPs from euchromatic regions of Chromosome 3. Scatter plots show relationships of principle components 1-8 where each marker represents an individual mosquito. Marker shape and colour denotes population. The bar chart shows the percentage of variance explained by each principal component.

**Figure S3.**
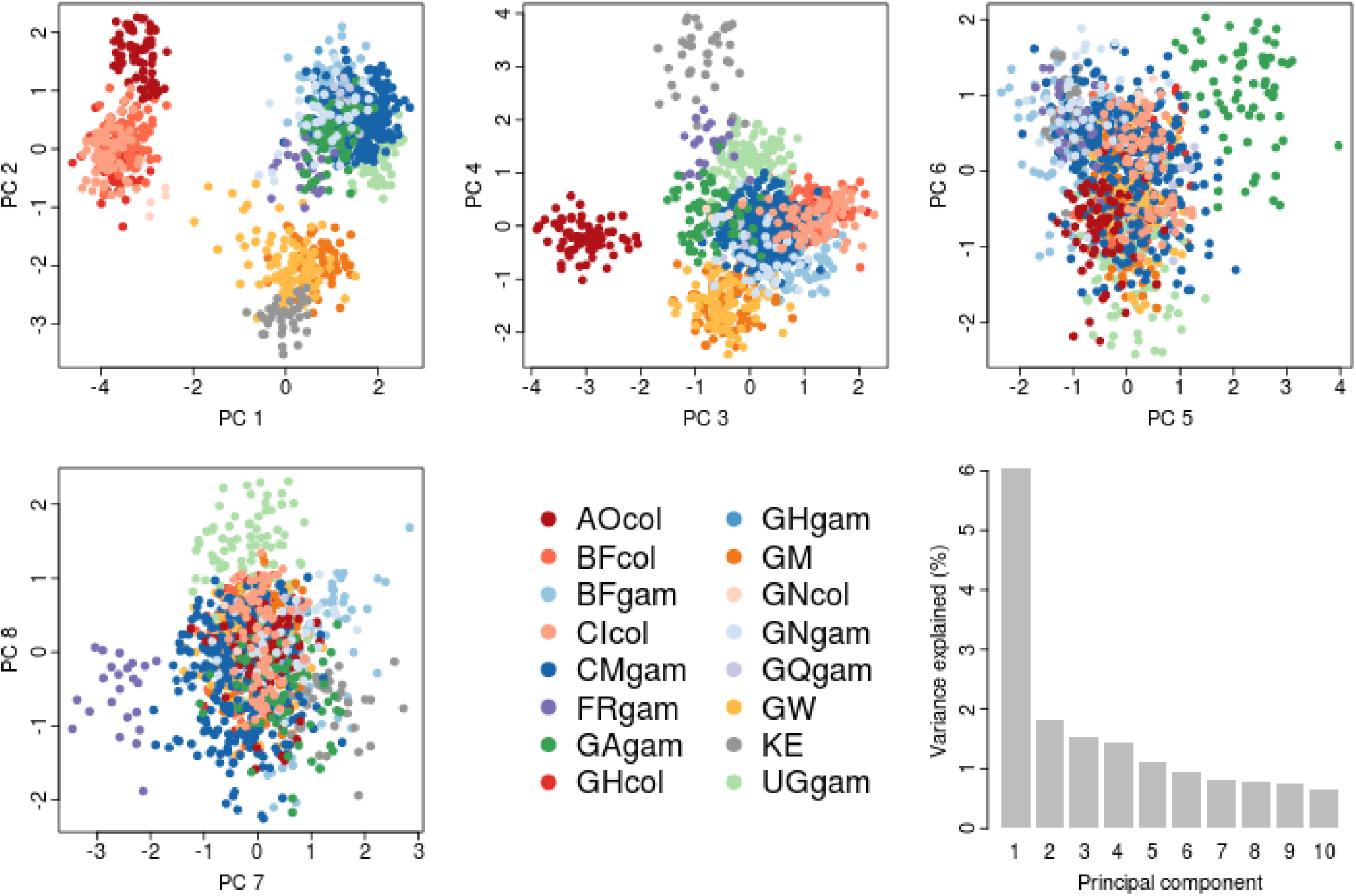
Principal component analysis of the 1,142 wild-caught mosquitoes using copy number variant calls. Bar chart shows the percentage of variance explained by each component.

**Figure S4.**
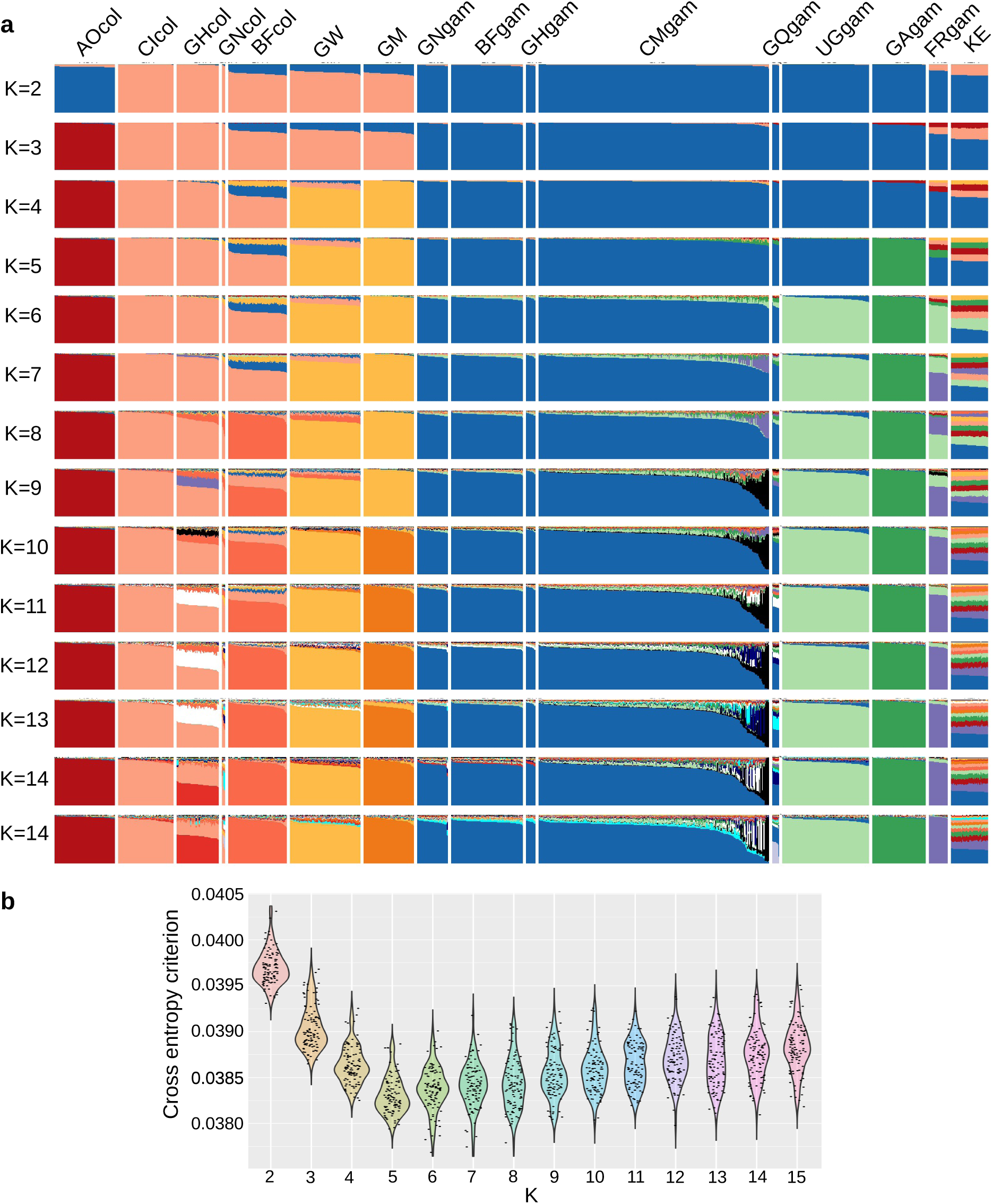
Analysis of population structure and admixture. **(a)** Each row shows results of modelling ancestry in sampled individuals assuming a given number *K* of ancestral populations [85]. Within each row, individual mosquitoes are represented as vertical bars, grouped according to sampling location and species, and coloured according to the proportion of the genome inherited from each ancestral population. **(b)** Cross-entropy criterion values obtained for each value of *K* ancestral populations, where lower values imply a better fit of the model to the data.

**Figure S5.**
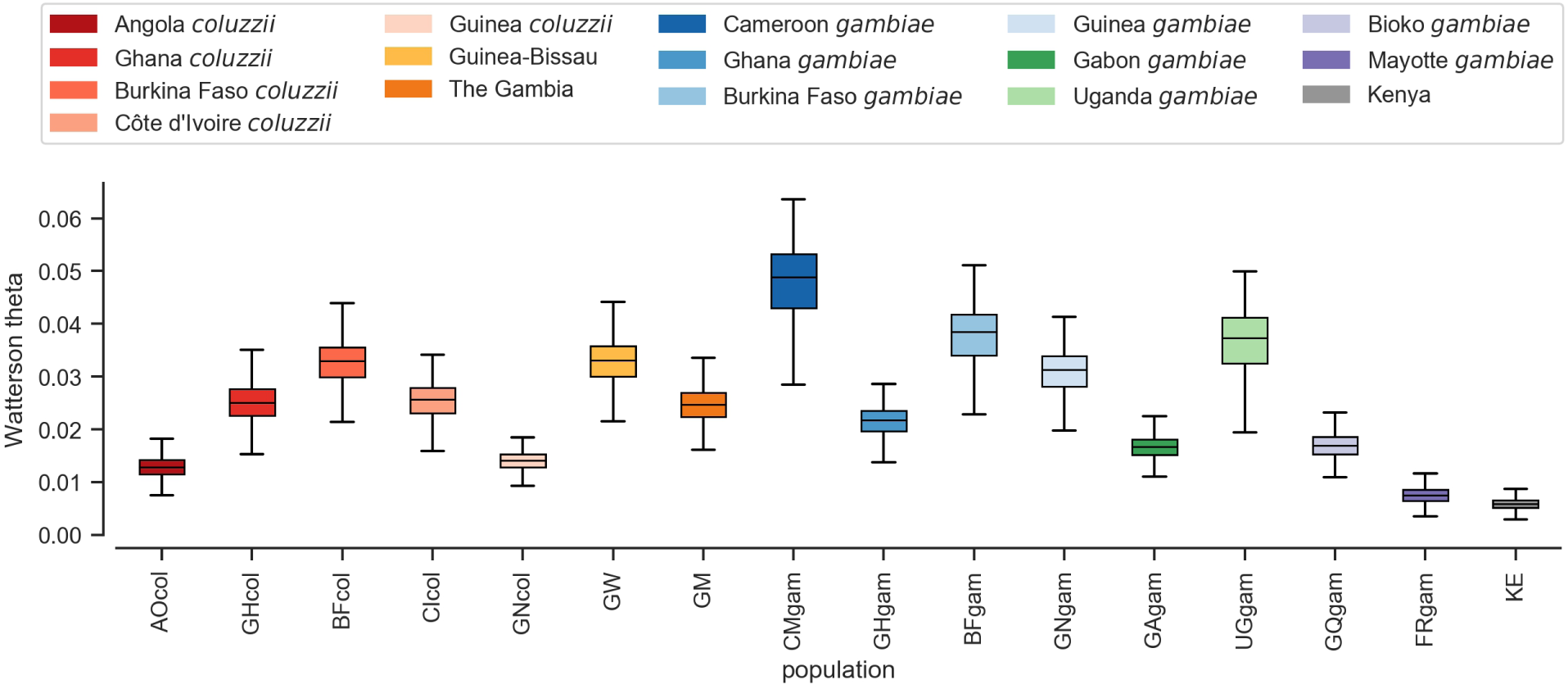
Watterson’s theta (*θ_W_*), the density of segregating sites, calculated in non-overlapping 20 kbp genomic windows using SNPs from euchromatic regions of Chromosome 3.

**Figure S6.**
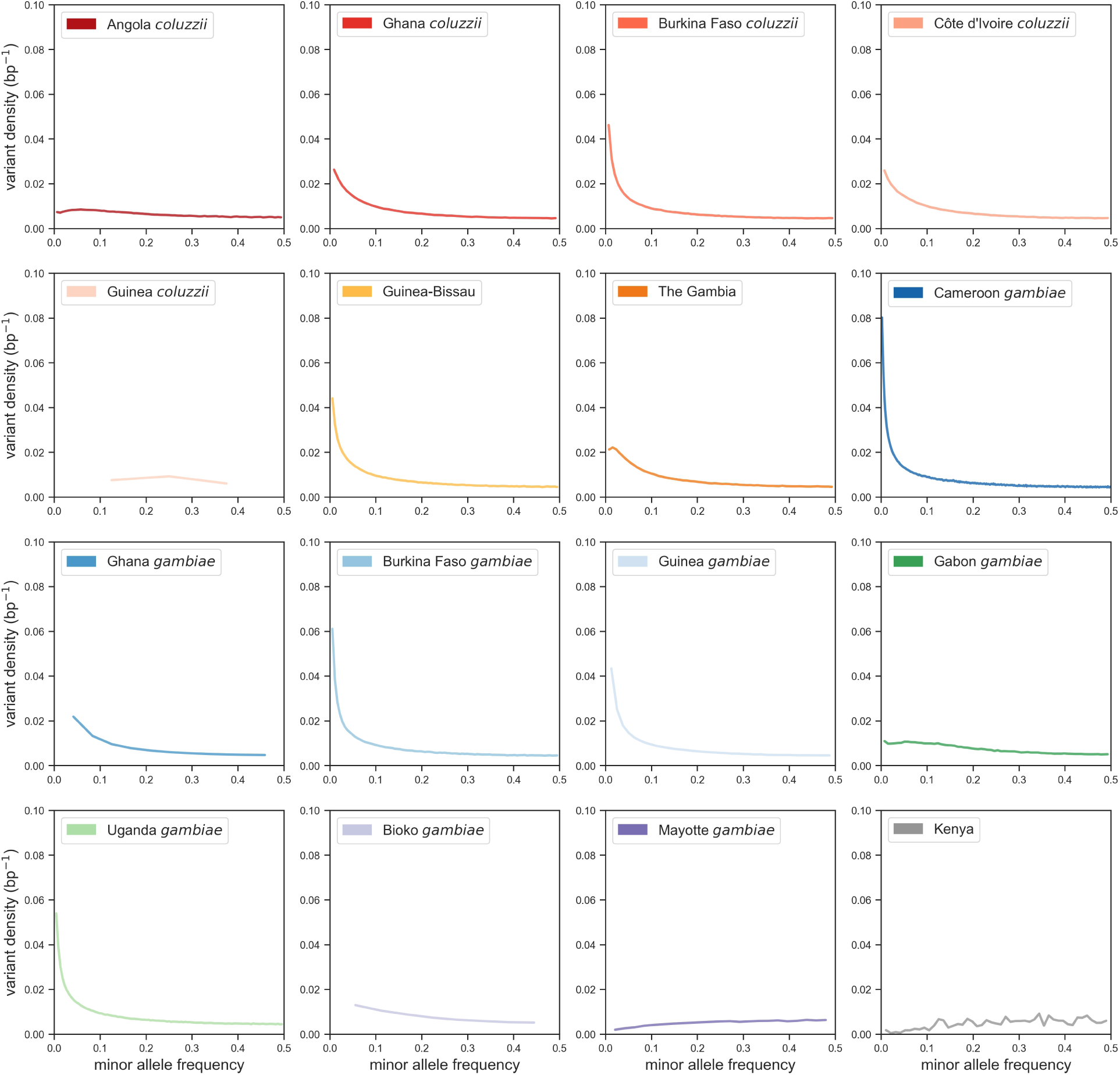
SNP density. Plots depict the distribution of allele frequencies (site frequency spectrum) for each population, scaled such that a population with constant size over time is expected to have a constant SNP density over all allele frequencies.

**Figure S7.**
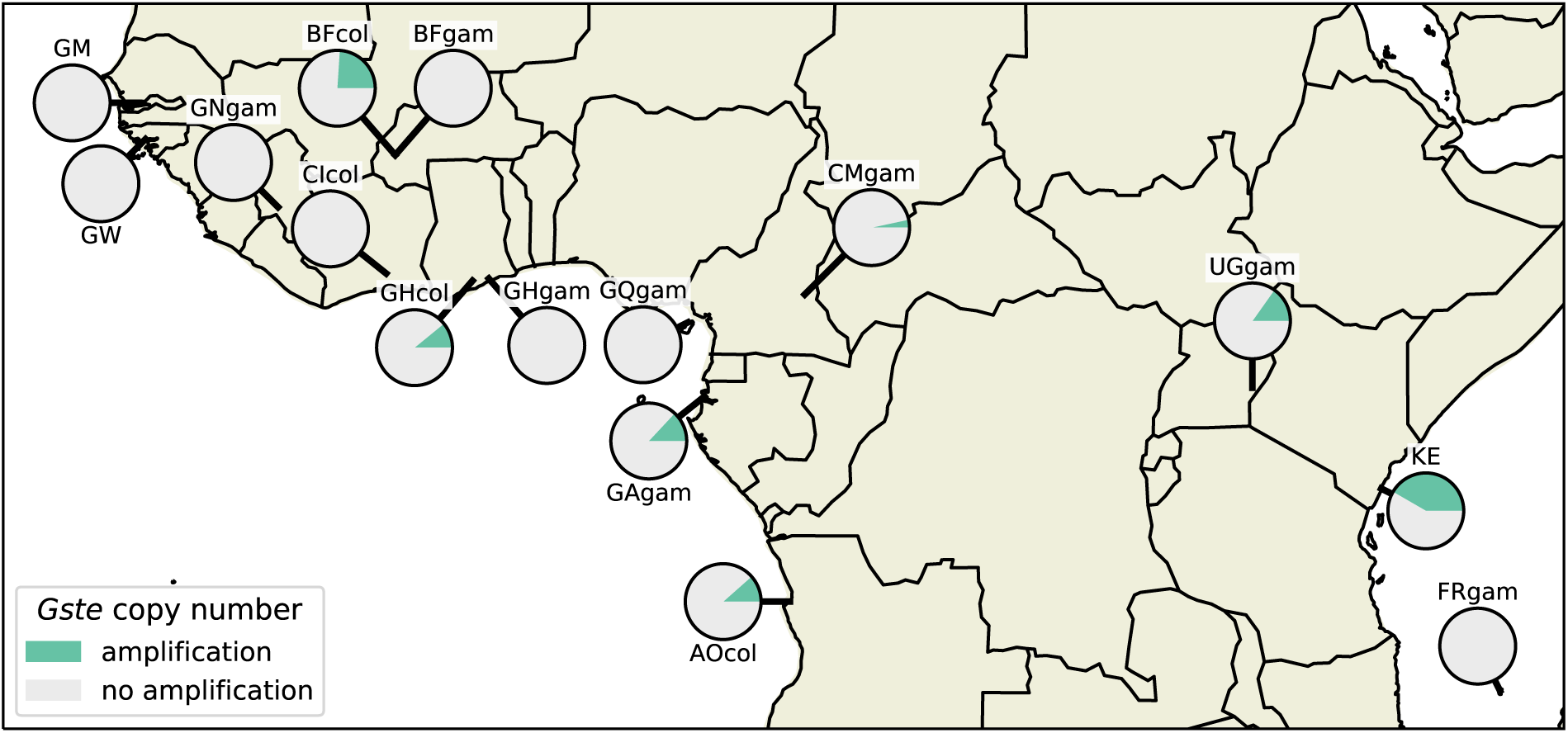
Prevalence of copy number amplifications at the *Gste* locus. Each pie shows the frequency of individuals from a given population carrying an amplification spanning at least one gene in the *Gste* gene cluster. The Guinea *An. coluzzii* population is omitted due to small sample size.

**Figure S8.**
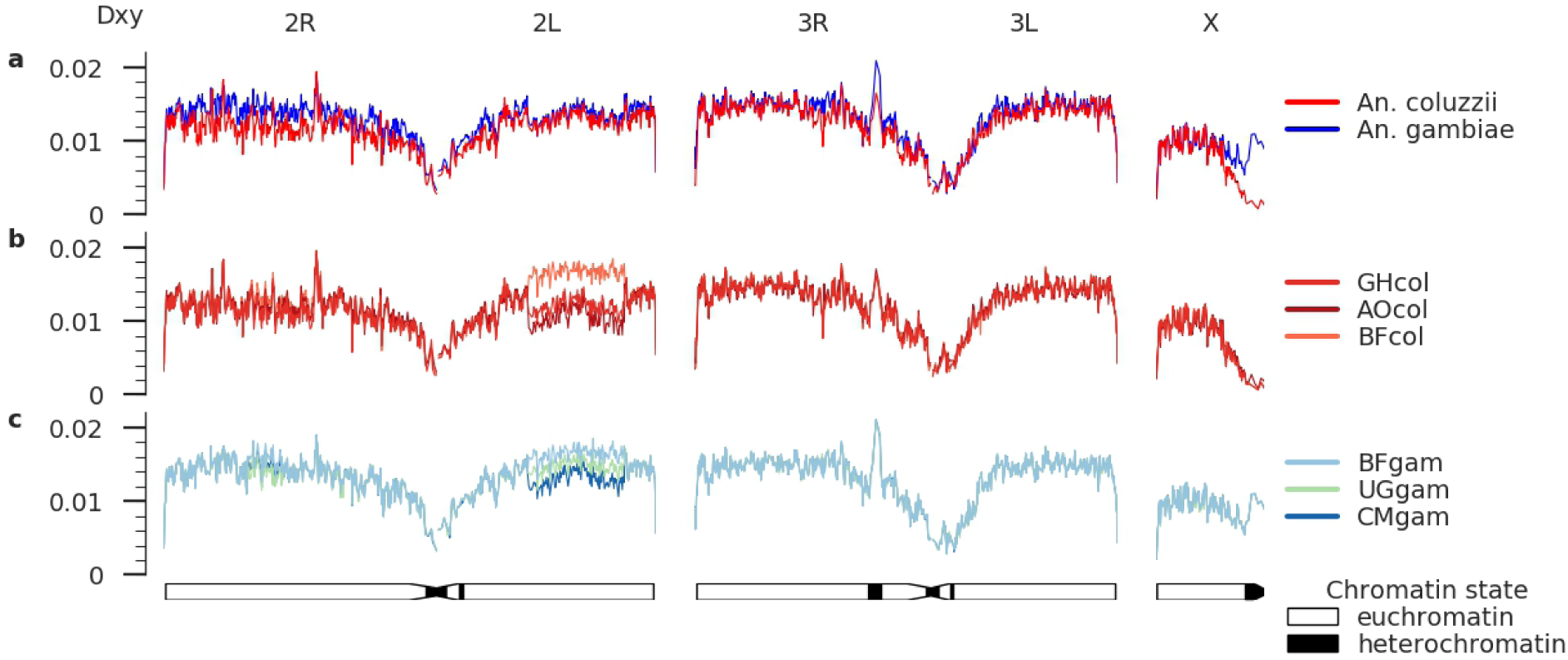
Divergence from the AgamP3 reference genome, calculated as *Dxy*, is largely similar for *An. coluzzii* and *An. gambiae*, with the exception of the centromere of the X chromosome (a). Comparing three populations of *An. coluzzii* (b) or *An. gambiae* (c) highlights the strong effect of the 2La chromosomal inversion on the accumulation of genetic variation.

**Table S1.** Ag1000G phase 2 sampling locations.

| Collection |  |  |  |  |  | Sample size |  |  |
| --- | --- | --- | --- | --- | --- | --- | --- | --- |
| Country | Location | Site | Year | Latitude | Longitude | Total | Female | Male |
| Angola | Luanda |  | 2009 | -8.821 | 13.291 | 78 | 78 | 0 |
| Burkina Faso | Bana |  | 2012 | 11.233 | -4.472 | 60 | 40 | 20 |
|  | Pala |  | 2012 | 11.150 | -4.235 | 56 | 48 | 8 |
|  | Souroukoudinga |  | 2012 | 11.235 | -4.535 | 51 | 51 | 0 |
| Cameroon | Daiguene |  | 2009 | 4.777 | 13.844 | 96 | 81 | 15 |
|  | Gado Badzere |  | 2009 | 5.747 | 14.442 | 73 | 58 | 15 |
|  | Mayos |  | 2009 | 4.341 | 13.558 | 105 | 91 | 14 |
|  | Zembe Borongo |  | 2009 | 5.747 | 14.442 | 23 | 23 | 0 |
| Cote d'Ivoire | Tiassale |  | 2012 | 5.898 | -4.823 | 71 | 71 | 0 |
| Equatorial Guinea | Bioko |  | 2002 | 3.700 | 8.700 | 9 | 9 | 0 |
| France | Mayotte | Bouyouni | 2011 | -12.738 | 45.142 | 1 | 1 | 0 |
|  |  | Combani | 2011 | -12.779 | 45.143 | 5 | 2 | 3 |
|  |  | Karihani Lake | 2011 | -12.797 | 45.122 | 3 | 3 | 0 |
|  |  | Mont Benara | 2011 | -12.857 | 45.155 | 2 | 1 | 1 |
|  |  | Mtsamboro Forest Reserve | 2011 | -12.703 | 45.081 | 1 | 1 | 0 |
|  |  | Mtsanga Charifou | 2011 | -12.991 | 45.156 | 8 | 3 | 5 |
|  |  | Sada | 2011 | -12.852 | 45.104 | 4 | 1 | 3 |
| Gabon | Libreville |  | 2000 | 0.384 | 9.455 | 69 | 69 | 0 |
| Gambia, The | Njabakunda | Kerr Birom Kardo | 2011 | 13.550 | -15.900 | 19 | 19 | 0 |
|  |  | Kerr Sama Kuma | 2011 | 13.550 | -15.900 | 8 | 8 | 0 |
|  |  | Maria Samba Nyado | 2011 | 13.550 | -15.900 | 18 | 18 | 0 |
|  |  | Sare Illo Buya | 2011 | 13.550 | -15.900 | 20 | 20 | 0 |
| Ghana | Koforidua |  | 2012 | 6.094 | -0.261 | 1 | 1 | 0 |
|  | Madina |  | 2012 | 5.668 | -0.219 | 24 | 24 | 0 |
|  | Takoradi |  | 2012 | 4.912 | -1.774 | 20 | 20 | 0 |
|  | Twifo Praso |  | 2012 | 5.609 | -1.549 | 22 | 22 | 0 |
| Guinea | Koraboh |  | 2012 | 9.250 | -9.917 | 22 | 22 | 0 |
|  | Koundara |  | 2012 | 8.500 | -9.417 | 22 | 22 | 0 |
| Guinea-Bissau | Antula |  | 2010 | 11.891 | -15.582 | 58 | 58 | 0 |
|  | Safim |  | 2010 | 11.957 | -15.649 | 33 | 33 | 0 |
| Kenya | Kilifi | Junju | 2012 | -3.862 | 39.745 | 16 | 16 | 0 |
|  |  | Mbogolo | 2012 | -3.635 | 39.858 | 32 | 32 | 0 |
| Uganda | Tororo | Nagongera | 2012 | 0.770 | 34.026 | 112 | 112 | 0 |

**Table S2.**
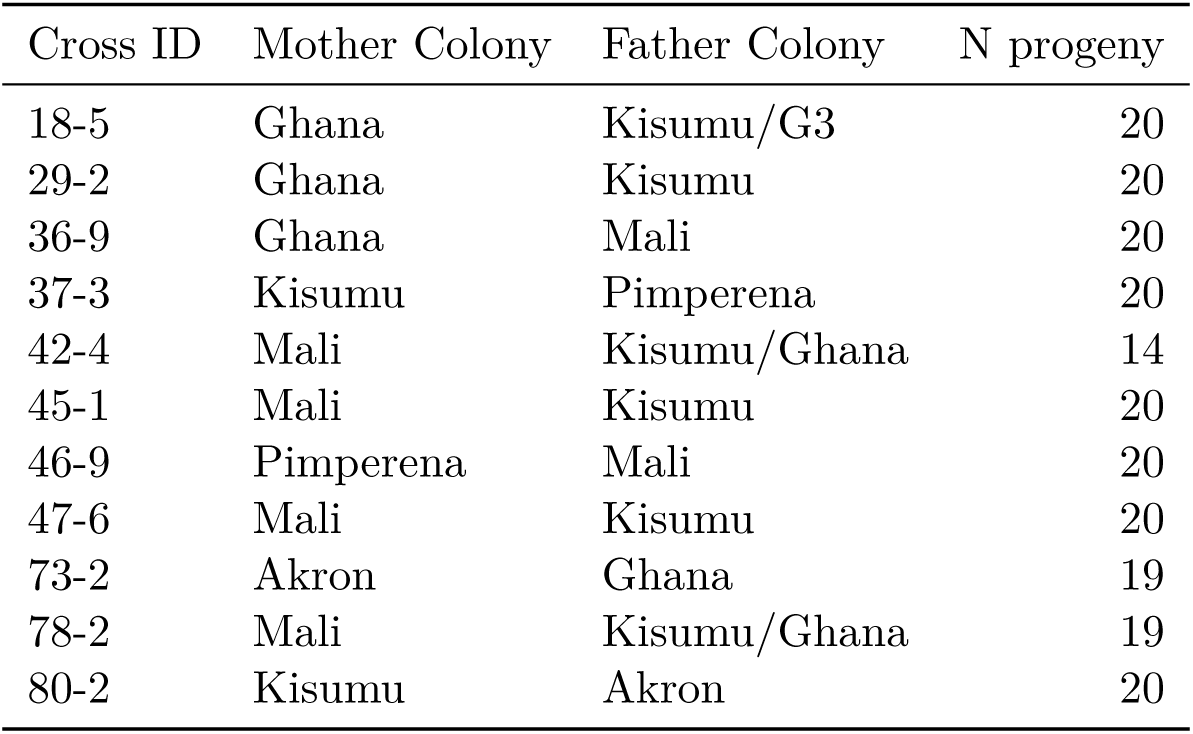
Colony crosses.

## Footnotes

1 https://www.malariagen.net/projects/ag1000g

2 https://www.malariagen.net

